# Quantitative Phosphoproteomic Analyses Identify STK11IP as a Lysosome-Specific Substrate of mTORC1 that Regulates Lysosomal Acidification

**DOI:** 10.1101/2021.08.18.456810

**Authors:** Zhenzhen Zi, Zhuzhen Zhang, Qiang Feng, Chiho Kim, Philipp E. Scherer, Jinming Gao, Beth Levine, Yonghao Yu

**Affiliations:** Department of Biochemistry, UT Southwestern Medical Center, Dallas, TX, 75390, USA; Touchstone Diabetes Center, UT Southwestern Medical Center, Dallas, TX, 75390, USA; Harold C. Simmons Comprehensive Cancer Center, University of Texas Southwestern Medical Center, Dallas, TX, 75390, USA; Howard Hughes Medical Institute and Center for Autophagy Research, University of Texas Southwestern Medical Center, Dallas, TX, 75390, USA

## Abstract

The evolutionarily conserved serine/threonine kinase mTORC1 is a central regulator of cell growth and proliferation. mTORC1 is activated on the lysosome surface. However, once mTORC1 is activated, it is unclear whether mTORC1 phosphorylates local lysosomal proteins to regulate specific aspects of lysosomal biology. Through cross-reference analyses of lysosome proteomic with mTORC1-regulated phosphoproteomic, we identified STK11IP as a novel lysosome-specific substrate of mTORC1. mTORC1 directly phosphorylates STK11IP at S404. Knockout of STK11IP led to a robust increase of autophagosome-lysosome fusion and autophagy flux. Dephosphorylation of STK11IP at S404 represses the role of STK11IP as an autophagy inhibitor. Mechanistically, STK11IP binds to V-ATPase, and regulates the activity of V-ATPase. Knockout of STK11IP protects mice from fasting and Methionine and Choline-Deficient Diet (MCD) diet induced fatty liver. Thus, our study demonstrates that STK11IP phosphorylation represents a novel mechanism for mTORC1 to regulate lysosomal acidification, and points to STK11IP as a promising therapeutic target for the amelioration of diseases with aberrant autophagy signaling.

---

The evolutionarily conserved Ser/Thr protein kinase mTORC1 (mammalian target of rapamycin complex 1) functions as the core catalytic component of pathways that regulates a variety of cellular anabolic processes ^1–3^. mTORC1 is activated on the surface of lysosome ^2, 4, 5^. Two necessary conditions are required to activate mTORC1. First, mTORC1 is recruited to the lysosome surface by the V-ATPase-Ragulator-Rag complex in response to amino acid and other nutrients; Second, mTORC1 is then activated by the lysosomal-bound GTPase, Rheb, in response to the growth factor-PI3K-AKT-TSC1/2 signaling pathway ^2, 6, 7^. Recent work has focused on identifying the mechanism by which mTORC1 translocates to the lysosome in response to different nutrients ^8–10^. Although the upstream modulation of mTORC1 activity is well addressed, the downstream signaling network of mTORC1 is poorly characterized. The best known substrates of mTORC1 are the ribosome protein S6 kinase (S6K) and the e1F4E-binding protein (4EBPs), both of which are known to regulate protein synthesis ^11^. In addition, mTORC1 regulates the synthesis of lipids and adipose tissue differentiation through SREBP and PPAR-γ ^12, 13^. Recently, using large-scale quantitative mass spectrometry experiments, we and others identified additional mTORC1 substrates and targets (for example, GRB10, IGFBP5, FOXK1 and SRPK2) that are involved in a number of critical cellular anabolic processes (e.g., glycolysis and lipid synthesis) ^14–17^.

Intriguingly, a number of recent studies showed that the lysosome is a key signaling hub that communicates, in a bi-directional manner, with mTORC1. Specifically, the lysosome not only controls the activation of mTORC1, but also receives signal from mTORC1 to regulate a number of important biological processes ^18^. For example, mTORC1 phosphorylates TFEB (Transcription factor EB), which is a master regulator of lysosome and autogenic gene expression, at S142 and S211. These two phosphorylation events cause the binding of TFEB to 14-3-3 proteins, and then the blockade of its translocation from the cytosol to the nucleus ^18, 19^. As a conserved cellular response to nutrient deprivation, mTORC1 also controls autophagy through inhibiting two key initiators of autophagosome formation, the ULK1 (Atg1) and the Atg13 proteins ^20, 21^.

## Integrative Analyses of the mTORC1-Regulated, Lysosomal Phosphoproteome

A key question is that once mTORC1 is activated on the lysosome surface, whether mTORC1 phosphorylates, and signals through local lysosomal proteins. Phosphoproteomic experiments at the organelle level pose a critical challenge, owing to the low stoichiometry of protein phosphorylation, and hence, the requirement for large amounts of starting material. To overcome this issue, we developed a two-step method. In the first step, we used mass spectrometry to define the lysosome proteome, and also the global mTORC1-regulated phosphoproteome. In the second step, we cross-reference analyzed the two datasets, and bioinformatically extracted the potential lysosome-specific mTORC1 substrates.

Using a recently reported strategy ^22, 23^, we generated HEK293T cells stably expressing LAMP1 that is fused with a C-terminal GFP and Flag^3X^ tags (LGF: LAMP1-GFP-Flag^3X^) (Fig. 1A). The isolated lysosomes showed superior purity, compared to those obtained using the traditional ultracentrifugation-based approach (Fig. S1A-C). From this lysosome-proteomic experiment (two biological replicate experiments were performed), we were able to identify a total of 232 proteins (Table S1). Gene Ontology (GO) analyses of these proteins showed that they were localized in vacuole and lysosome. These proteins were highly enriched with biological processes including vesical-mediated transport, endocytosis and ion transport, all of which are known to be connected to lysosome biology (Fig. S1D-F) ^4, 24, 25^. We interrogated the list, and found that the identified proteins included both well-known lysosomal proteins, including LAMP1 and LAMP2; the Ragulator complex proteins; the V-ATPase complex proteins; hydrolases; ion transporters; trafficking and fusion machinery proteins (Fig. S1G) ^24^, as well as several recently reported lysosomal proteins, including SLC38A9 ^26, 27^ and aldolase A ^28^ (Fig. S1G).

**Fig. 1.**
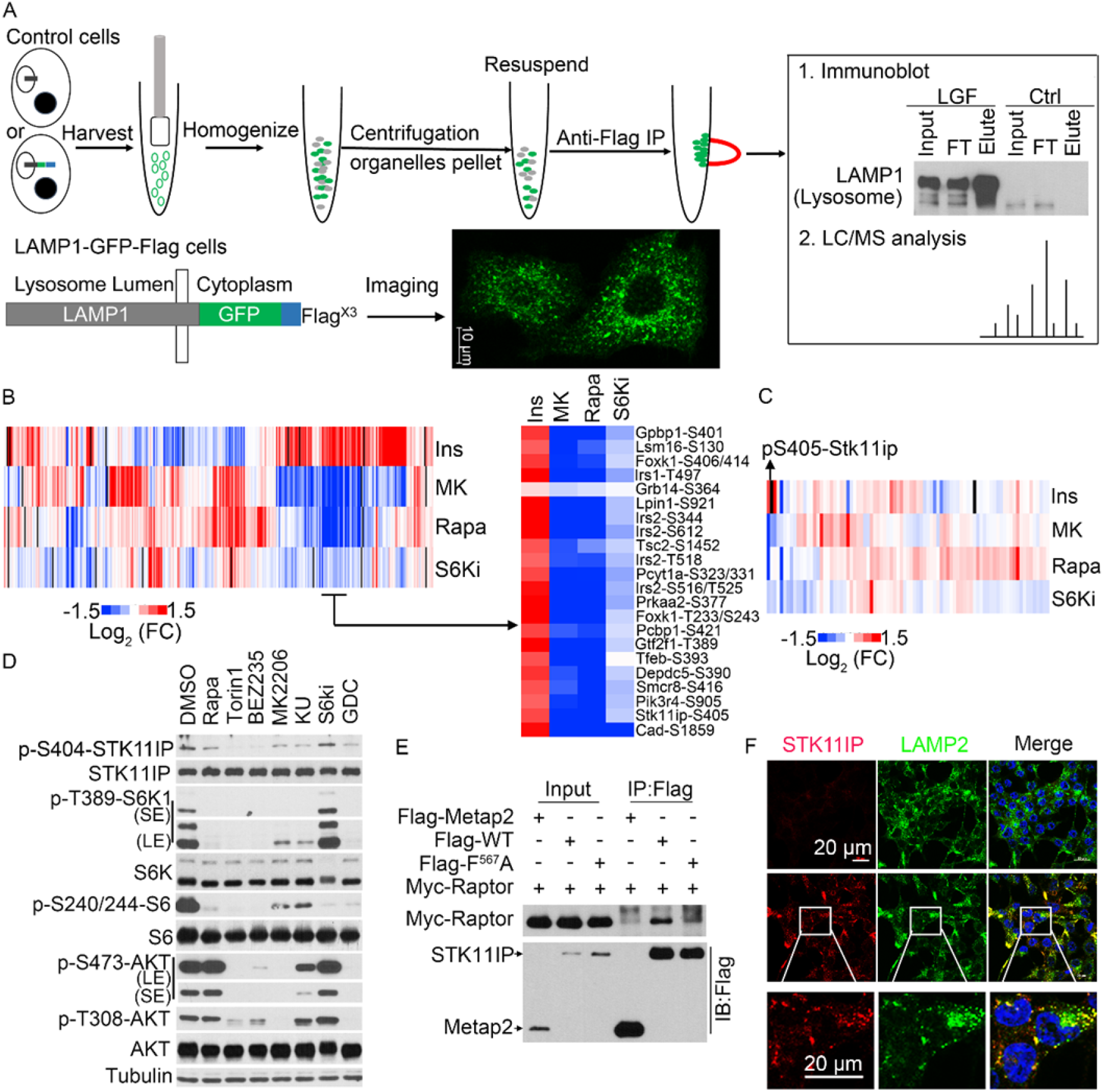
Integrative analyses of the mTORC1-regulated lysosomal phosphoproteome. (A) Schematic of the workflow for the Lysosome-IP method. Control and LAMP1-GFP-Flag stably expressing HEK293T cells were harvested and homogenized. Flag-tagged lysosomes were isolated with the anti-Flag magnetic beads and were subjected to WB or LC-MS/MS analyses. Confocal microscopy analyses were performed to validate the localization of the LAMP1-GFP signal. Scales bar, 10 μm. (B) Heat map representation of the phosphopeptides identified in the various screens in the hepatic phosphoproteomic. FC, fold change. Ins (Insulin, 30 min); MK (MK2206, Akt inhibitor, 2 hour); Rapa (rapamycin, mTORC1 inhibitor, 2 hour); S6Ki (S6 kinase inhibitor, 2 hour). (C) Heat map representation of the phosphorylation changes of the peptides commonly identified in the lysosome proteomic and hepatic phosphoproteomic. FC, fold change, pS405-Stk11ip in the rat proteins (pS404 in the human proteins). (D) Phosphorylation of STK11IP at S404 is blocked by mTOR inhibitors in HEK293T cells for 4 hours. DMSO treatment was used as the control; Rapamycin (Rapa, 20 nM, an mTORC1 inhibitor); Torin1 (2mM, an mTOR kinase inhibitor); BEZ235 (1 μM, a PI3K/mTORC1 dual inhibitor); MK2206 (1 μM, an Akt inhibitor); KU-0063794 (KU, 1 μM, an mTOR kinase inhibitor); PF-4708671 (S6Ki, 10 μM, an S6K inhibitor); GDC-0941 (1 μM, an PI3K inhibitor). (E) STK11IP interacts with Raptor. Flag-tagged STK11IP (WT and F^567^A mutant) or Flag-Metap2 (as the control) was co-transfected with Myc-Raptor into HEK293T cells. The lysates were subjected to immunoprecipitation using the anti-Flag beads. Raptor levels were probed using an antibody against the Myc-tag. (F) Immunofluorescence analysis of STK11IP and LAMP2 in the STK11IP wild type or knockout cells. Scales bar, 20 μm.

Using a reductive dimethylation-based quantitative mass spectrometry approach, we recently characterized the hepatic phosphoproteome (in primary rat hepatocytes) (Table S2) that is controlled by insulin, as well as its key downstream kinases, including Akt, mTORC1 and S6K ^16^. Cross-reference analyses showed that a total of 45 proteins were commonly identified between the hepatic phosphoproteomic datasets and the lysosomal proteomic results (Fig. 1B, 1C S1H and Table S3). We then extracted, from the hepatic phosphoproteomic datasets, the proteins that are potential mTORC1 substrates, i.e., those proteins with decreased phosphorylation upon rapamycin treatment, and unchanged phosphorylation upon the treatment of an S6K inhibitor. Among these proteins, only STK11IP was also identified in the lysosomal proteomic experiment (Fig. 1B, and 1C).

## STK11IP is a Lysosome-specific Substrate of mTORC1

Phosphorylation of STK11IP (S405 in the rat / mouse protein and S404 in the human protein; the phosphorylation site in the human protein will be used hereafter) decreased markedly (~6-fold) after rapamycin treatment (Fig. S1I). This phosphorylation site is conserved and positioned in a sequence (i.e., LDPS^404^PAG) that is consistent with known mTORC1 phosphorylation motifs (i.e., a Pro residue at the +1 position) ^29^ (Fig. S1J). We found that phosphorylation pattern of STK11IP (pS404) was similar to other known mTORC1 substrates, including TFEB (pS393) and ULK1 (pS757), but was different from known substrates of S6K (an AGC kinase that prefers basic residues in the vicinity of the phosphorylation site), including Cad (pS1859) and rpS6 (pS236/240) (Fig. S1K) ^21, 30, 31^. Collectively, these results suggested STK11IP (pS404) as a direct substrate of mTORC1.

We generated a phospho-specific antibody against STK11IP/pS404 (Fig. S2A and S2B) and found that pS404 was potently inhibited by Rapamycin, and mTOR kinase inhibitors (e.g., Torin1 and BEZ235). This phosphorylation site, however, was refractory to inhibitors of S6K (Fig. 1D), JNK1 or ERK (Fig. S2C). pS404-STK11IP levels were also dramatically downregulated during amino acid starvation (Fig. S2D). Moreover, mTOR was able to phosphorylate STK11IP (pS404) in an *in vitro* kinase assay (Fig. S2E). We found that STK11IP interacted with Raptor, mTOR and PRAS40, but not Rictor (Fig. 1E and S2F-S2H). Finally, inspection of the STK11IP sequence suggested that this protein contained a potential TOS motif (F^567^EVEL) (TOS motif refers to the TOR signaling motif ^32^). Indeed, the mutation of the key residue (F567) in this TOS motif (F567A) abolished the binding between STK11IP and Raptor (Fig. 1E). These biochemical results further demonstrated that STK11IP is a direct substrate of mTORC1.

We then confirmed that STK11IP existed in the lysosome fraction using both immunoprecipitation and ultracentrifugation approaches (Fig. S1A and S1B). This was further validated by inverse immunoprecipitation experiments using the STK11IP-GFP-Flag (SGF) stable cells, which showed that only lysosome could be immunoprecipitated (Fig. S2I). Co-immunostaining assays showed that endogenous STK11IP was co-localized with LAMP2 (a lysosome marker that is localized on the lysosome membrane), but not with Tom20 (a mitochondria marker) (Fig. 1F and Fig. S2J). Finally, we found that STK11IP was distributed in the membrane fraction using subcellular fractionation experiments (Fig. S2K). Taken together, these results suggested that STK11IP is an mTORC1 substrate that is resident on the lysosomal membrane.

## STK11IP Deficiency Promotes Autophagy

STK11IP is a poorly characterized protein ^33, 34^. A number of mTORC1 substrates are known to regulate its activation through feedback mechanisms ^3, 35^. However, we did not observe any differences in mTORC1 activity between control and STK11IP knockout (KO) primary MEF cells, as shown by their similar pS6K and pULK1 levels (Fig. S3A). We used the CRISPR-Cas9 system to generate the STK11IP KO cells, and reconstituted these cells with WT-, S404A- or S404D-STK11IP. We performed amino acid starvation experiments and found no differences in mTORC1 activities in these cells (Fig. S3B). These experiments suggest that the activation of mTORC1 is not affected by STK11IP, or its mTORC1-mediated phosphorylation.

We next investigated whether STK11IP could mediate certain biological processes downstream of mTORC1. In particular, mTORC1 is critically involved in controlling autophagy, via several of its downstream target proteins (e.g., TFEB and ULK1) ^21, 24^. Here, we found that the downregulation of pS404-STK11IP occurred together with the upregulation of autophagy levels, as shown by the increased conversion of LC3-I to LC3-II (LC3II/LC3I ratio) during mTORC1 inhibition (Fig. S2D). These results suggest that STK11IP may act as a downstream target of mTORC1 to regulate autophagy. To directly test this hypothesis, we knocked out STK11IP in HEK293T cells using the CRISPR-Cas9 technology. Compared to control cells, LC3II, the marker for autophagosome numbers, was markedly increased in STK11IP KO cells (Fig. 2A and 2B). Besides, there were more LC3 puncta (endogenous LC3) or GFP-LC3 puncta (using GFP-LC3 stable cells) in STK11IP KO cells compared to those in the control cells (Fig. 2C and 2D). STK11IP KO also leads to a profound increase in LC3 or GFP-LC3 puncta numbers under chloroquine (CQ) treatment compared to control conditions (Fig. 2C and 2D).

**Fig. 2.**
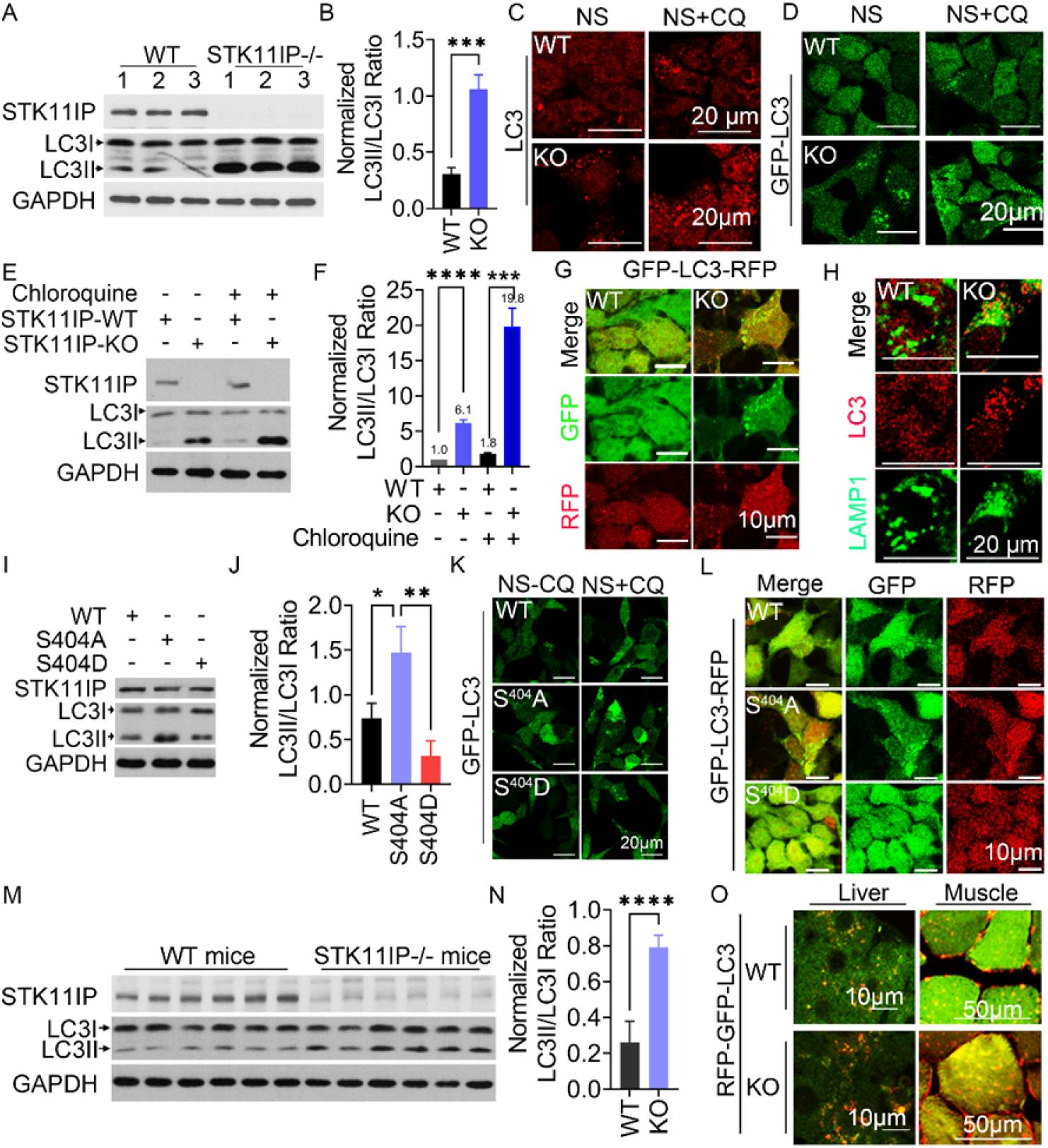
STK11IP Deletion Promotes Autophagy in Cells or Mice under Endurance Exercise Test. (A-B) Immunoblot analyses (A) and quantification (B) of the LC3II and LC3I levels in STK11IP WT (wild type) or KO (Knock out) cells; (n=3 per group), t test. Results represent mean ± SEM. (C-D) STK11IP KO (knockout) cells have more LC3 puncta (endogenous LC3) (C) and GFP-LC3 puncta (using GFP-LC3 stable cells) (D) than WT (wild type) cells under the indicated conditions, NS (non-starvation), CQ treatment for 2 hours. Scales bars, 20 μm. (E-F) STK11IP knock out leads to increased LC3 turnover. WT and STK11IP KO cells were cultured in complete media. When indicated, chloroquine treatment was performed 20 μM for 2 hours. Immunoblot analyses (E) and quantification of the LC3II/LC3I ratio (F); (n=3); t test. Results represent mean ± SEM. (G) STK11IP Knockout leads to increased autophagic flux. The GFP-LC3-RFP probe was stably expressed in either WT or STK11IP KO cells. The cells were cultured in complete media. Scales bars, 10 μm. (H) STK11IP knock out leads to more co-localization of LC3 puncta and LAMP1 puncta, which indicate enhanced formation of autolysosome. Co-immunostaining of LAMP1 (Green signal) and LC3 (Red signal) was performed using STK11IP wild-type and knock-out cells. The cells were cultured in complete media. Scales bars, 20 μm. (I-J) Immunoblot analyses (I) and quantification (J) of the LC3II and LC3I levels in WT, S404A and S404D reconstitute cells; (n=3), t test. Results represent mean ± SEM. The cells were cultured in complete media. (K) STK11IP-S404A mutant leads to the increased formation of GFP-LC3 puncta. Immunofluorescence of GFP-LC3 in the WT, S404A or S404D reconstituted cells under indicated conditions. When indicated, chloroquine treatment was performed at 20 μM for 2 hours, NS (non-starvation). Scales bars, 20 μm. (L) Immunofluorescence analyses of the GFP-LC3-RFP signal in WT, S404A and S404D reconstituted cells. Cells were cultured in complete media. Scales bars, 10 μm. (M-N) STK11IP KO leads to increased autophagic flux in mice. Western blot analysis (M) and quantification (N) of LC3-II/LC3-I level in soleus muscle tissues from WT (wild type) and STK11IP KO (knockout) mice, after the endurance exercise test; (n=6 per group), t test. Results represent mean ± SEM. (O) Representative images of the mRFP-GFP-LC3 puncta in the liver (Left) and soleus muscle (Right) from STK11IP WT (wild type) and KO (knockout) mice after endurance exercise test that stably express mRFP-GFP-LC3 (tfLC3). Scales bars, 10 μm for liver and 50 μm for soleus muscle.

Both autophagy activation and decreased autophagic degradation will lead to the increase of the autophagosome number ^36^. The LC3 turnover assay measures the autophagic flux, which is therefore a more reliable indicator of the autophagic activity ^36^. We performed the LC3 turnover assay to compare the autophagy flux between WT and STK11IP KO cells. Our results showed that STK11IP knockout increases autophagy flux by around 2-fold (LC3II/LC3I ratio with CQ versus without CQ in WT and KO cells) (Fig. 2E, 2F and Fig. S3C). Another commonly used assay for autophagic flux measurement is based on a fluorescent probe, GFP-LC3-RFP ^37^. Because of the different stability between GFP and RFP signal under acidic conditions, the GFP/RFP ratio reflects the autophagic flux ^37, 38^. We then applied the fluorescent probe assay and found that STK11IP KO cells displayed a lower GFP/RFP ratio (stronger yellow color) and more GFP-LC3 puncta, compared to control cells (Fig. 2G, S3D and S3E), indicating the enhanced autophagy flux in the STK11IP KO cells. More specifically, we found that autophagosome-lysosome fusion was enhanced in STK11IP knockout cells (Fig. 2H and S3F). We observed increased co-localization of LC3 and LAMP1 in STK11IP-/- cells compared to wild type cells, suggesting that there is an increased number of autolysosomes in STK11IP-/- cells under both basal and amino acid-starvation conditions. Thus, the enhanced autophagy flux could be the result of increased autophagosome-lysosome fusion.

To investigate the role of the mTORC1-mediated phosphorylation of STK11IP in autophagy, we expressed the STK11IP WT, S404A or S404D mutant in STK11IP KO cells. Both LC3II levels and GFP-LC3 puncta numbers were increased in cells that stably expressed the S404A-STK11IP mutant, compared to WT and S404D mutants (Fig. 2I, 2J and 2K). To measure the autophagy flux, we also expressed the GFP-LC3-RFP probe in the cells expressing the STK11IP WT, S404A or S404D mutant. Similar to STK11IP KO cells, S404A mutants showed a higher autophagy flux, as demonstrated by the reduced GFP/RFP signal and more GFP-LC3 puncta (Fig. 2L). Collectively, our results suggest that dephosphorylation of STK11IP at S404 or STK11IP deficiency promotes autophagy flux.

We next utilized the STK11IP KO mice to determine whether STK11IP regulates autophagy *in vivo*. One of the most potent physiological inducers of autophagy is exercise ^39, 40^. To test whether STK11IP can regulate exercise-induced autophagy *in vivo*, WT and STK11IP KO littermates were subject to the endurance exercise test (Fig. S3G-S3K). We found that the STK11IP KO mice exhibited much higher levels of exercise-induced LC3-II conversion in the soleus muscle (Fig. 2M and 2N). The pS404-STK11IP level was decreased upon endurance treadmill exercise test (Fig. S3L). We also generated a mouse line by crossing the STK11I KO mice with the TfLC3 (mRFP-GFP-LC3) mice ^41^. We then assessed the number of mRFP-GFP-LC3 puncta in the liver and soleus muscle after the endurance exercise test. Remarkably, STK11IP KO mice displayed reduced GFP/RFP ratios and enhanced mRFP-GFP-LC3 puncta in both liver and soleus, which indicated a dramatic upregulation of autophagy flux in these mice (Fig. 2O and Fig. S3M-S3N). In summary, STK11IP KO promotes autophagy flux *in vivo*.

## STK11IP Binds to V-ATPase to Regulate Lysosomal Acidification

We investigated the potential molecular mechanism by which STK11IP regulates autophagy. Autophagy is a multi-step process, which includes, the initiation, autophagosome formation, and the autophagosome-lysosome fusion ^36, 42^. However, we did not observe differences in the level of lysosome biogenesis and lysosome morphologies in control vs. STK11IP KO cells (Fig. S4A and S4B).

As mentioned above, more co-localization of LC3 and LAMP1 was observed in the STK11IP KO cells, suggesting that STK11IP could regulate autophagosome and lysosome fusion (Fig. 2H and Fig. S3F). Previous reports showed that autophagosome-lysosome fusion could be regulated by the pH of cellular acidic compartments ^43^. It is therefore likely that the STK11IP could regulate lysosomal acidification (Fig. 2G, 2L). Intriguingly, mTORC1 inactivation by Torin1 treatment or amino acid starvation could lead to decrease of pH value (Fig. S4C) and activation of autophagy (Fig. S2D) ^36^. These results raise the hypothesis that mTORC1 could regulate lysosomal acidification via an STK11IP-dependent mechanism. Indeed, we observed that the lysosomes were more acidic in STK11IP KO cells relative to the controls (Fig. 3A and S4D). Furthermore, STK11IP KO cells reconstituted with the STK11IP S404A mutant also showed more acidic lysosomes, compared to those derived from the cells expressing STK11IP WT or the S404D mutant (Fig. 3B). We also used a ratiometric probe, LysoSensor yellow/blue dye DND-160, to measure the lysosomal pH value (based on the ratio of yellow/blue signals) ^44, 45^. The results showed that the pH value is significantly lower in STK11IP KO cells and STK11IP S404A expressing cells (Fig. 3C, 3D, S4E and S4F), compared to their control cells. Recently, a series of ultra-pH sensitive (UPS) nanotransistor probes were reported, which display much sharper on/off pH responses ^46, 47^. We observed that the lysosomal pH value was lower than 5.4 in both the STK11IP KO cells and WT/S404A/S404D reconstituted cells. However, the activated TMR signal can only be detected in the STK11IP KO and S404A reconstituted cells, when the pH 4.5 nanoprobe was applied (Fig. 3E and 3F). This result is also consistent with the pH value measured using the lysoSensor yellow/blue dye DND-160 method (Fig. 3C and 3D).

**Fig. 3.**
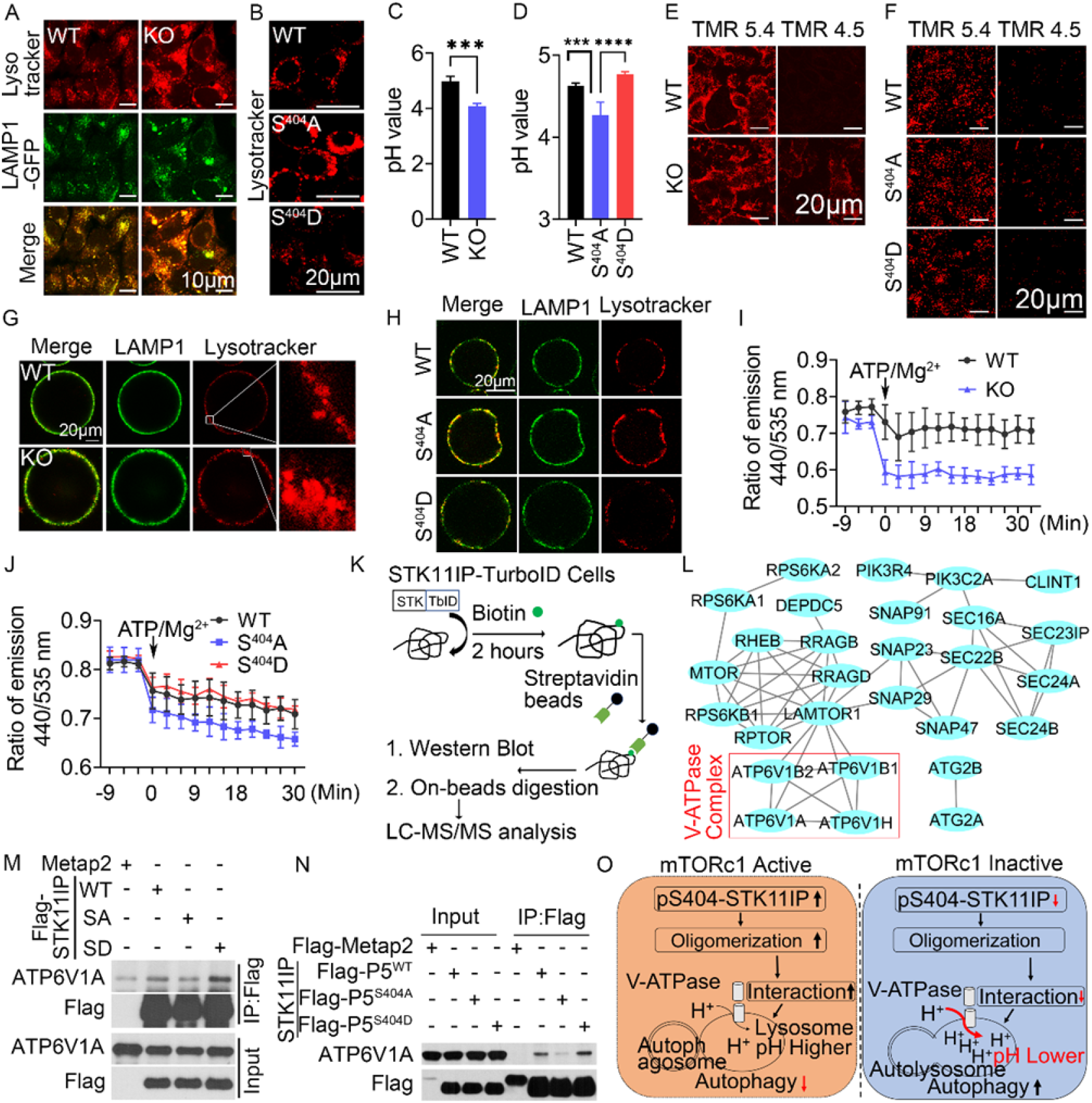
STK11IP Regulate Lysosomal Acidification. (A) STK11IP knock out leads to increased lysosomal acidity. WT (wild type) or KO (knock out) cells were stained with the Lysotracker dye and subject to immunofluorescence analyses. Scales bar, 10 μm (A). (B) STK11IP-S404A mutant leads to increased lysosomal acidity compared to WT and S404D reconstituted cells. The WT, S404A or S404D reconstituted cells were stained with the Lysotracker dye and analyzed by immunofluorescence analyses. Scales bar, 20 μm. (C) pH values were measured in the WT and STK11IP KO cells. Cells were cultured in the normal media and stained with the LysoSensor Yellow/Blue DND-160 dye. The fluorescence signal was measured using a microplate reader (n=4 per group); t test. Results represent mean ± SEM. (D) pH values were measured in WT, S404A and S404D reconstituted cells, same to (C). (n=4 per group); t test. Results represent mean ± SEM. (E-F) pH imaging of the acidic organelles using the TMR (tetramethyl rhodamine)-HyUPS nanoprobes in WT and KO (knock out) live cells (E) or WT/S404A/S404D reconstituted cells (F). Cells were incubated with 25 μg mL^-1^ of the TMR-HyUPS nanoprobe at 37 °C for 15 min and imaged by confocal microscopy. Scale bars, 20 μm. (G) STK11IP knock out leads to increased lysosomal acidity in the isolated lysosomes. Lysotracker staining for the lysosomes, which were immunoprecipitated from WT/KO (knock out) HEK293T cells. Scale bars, 20 μm. (H) STK11IP-S404A mutant leads to increased lysosomal acidity in the isolated lysosomes. Lysotracker staining for the lysosomes, which were immunoprecipitated from WT/S404A/S404D reconstituted HEK293T cells. Scale bars, 20 μm. (I) V-ATPase activity is enhanced in STK11IP knock out cells compared to WT cells. The activity of V-ATPase was measured using the Lysosensor Yellow/Blue DND160 probe in the isolated lysosomes (from WT and STK11IP KO HEK293T cells). The pH gradient across the lysosomal membrane was intentionally dissipated during the course of isolation by FCCP. Where indicated, lysosomal re-acidification was initiated by addition of ATP/Mg^2+^. The light emitted at 440 and 535 nm was measured with microplate reader and ratio of 440/535 was calculated, (n=6 per group), Results represent mean ± SEM. (J) V-ATPase activity is enhanced in S404A reconstituted cells compared to WT and S404D cells. The procedure is same to (I). (K-L) Schematic of the workflow for the STK11IP-TurboID experiment (K). Functional association analyses of the STK11IP-interacting proteins identified from the TurboID MS experiment (L). The protein-protein interaction network was generated using the STRING database and viewed by CytoScape. V-ATPase complex was highlighted in the red box. (M) The STK11IP S404A (SA) mutant interacts more weakly with ATP6V1A, compared to STK11IP WT or the S404D (SD) mutant. (N) The S404A mutant-P5 domain of STK11IP mutant interacts more weakly with ATP6V1A, compared to WT and the S404D P5 mutant. P5: STK11IP (360-770); Metap2 was used as the negative control. (O) The scheme of STK11IP in regulating lysosomal acidification and autophagy.

We further explored the potential mechanism by which STK11IP regulates lysosomal acidification. Lysosomes maintain their pH gradients through the proton-pumping V-type ATPase ^48, 49^. We therefore asked whether STK11IP has an impact on the V-ATPase activity. We isolated the lysosomes from WT and STK11IP KO cells, or STK11IP-WT, S404A and S404D reconstituted cells, and checked the V-ATPase activity using an acidification assay ^50^. We found that the activity of V-ATPase was enhanced in the lysosomes isolated from STK11IP KO and STK11IP S404A cells compared to their control cells (Fig. 3G, 3H, 3I and 3J).

Next we sought to understand how STK11IP controls the activity of V-ATPase. First, we performed quantitative proteomic analyses (using the tandem mass tag, TMT labeling approach) of STK11IP WT and KO cells. Through these experiments, we found that STK11IP KO has no effect on the expression of the various components of V-ATPase (Fig. S5A, and table S4). Next, we examined whether STK11IP can interact with the V-ATPase complex. We used the TurboID system to identify the STK11IP-interacting proteins ^51^ (Fig. 3K). We observed that many of these biotinylated proteins were localized on the lysosome and were involved in mTORC1 signaling, including the Rag-GTPase A/D, RAPTOR, mTOR and LAMTOR1. Although STK11IP has been reported to bind LKB1 and regulate TGF-beta signaling ^33, 34^, we did not identify LKB1 in our TuroID-MS dataset, potentially due to the different experiment conditions. Notably, we also identified several components of the V-ATPase complex, including ATP6V1B, ATP6V1A and ATP6V1H (Fig. 3L, S5B and table S5). The interaction between STK11IP and the various subunits of the V-ATPase complex was further validated by co-immunoprecipitation (Fig. S5C and S5D). Through domain mapping experiments, we further identified that the domain of amino acid 360 to amino acid 770 of STK11IP mediated this interaction between STK11IP and ATP6V1A (Fig. S5E and S5F).

Interestingly, we found that compared to the WT protein, the S404A mutant formed a much weaker interaction with the V-ATPase components (ATP6V1A and ATP6V1B) (Fig. 3M, and S5G). Compared to the WT STK11IP domain of amino acid 360 to amino acid 770, the same domain bearing the S404A mutation displayed a weaker interaction with ATP6V1A (Fig. 3N). These data indicated that mTORC1-mediated phosphorylation of STK11IP at S404 is essential for its interaction with the V-ATPase complex. Taken together, mTORC1 signaling regulates the phosphorylation of STK11IP at S404, and de-phosphorylation of S404 disrupts its interactions with the V-ATPase. The de-phosphorylation of STK11IP results in increased V-ATPase activity, more acidic lysosomes, and hence enhanced autophagy flux (Fig. 3O).

## STK11IP Deficiency Protects against Fasting- and MCD-Induced Fatty Liver

Autophagy occurs in virtually all cells to maintain cellular homeostasis, including modulation of nutrient balances ^42, 52^. To investigate the physiological function of STK11IP in mice, we first measured the expression profile of STK11IP in mouse tissues. Results from qRT-PCR and immunoblot analyses showed that *Stk11ip* is widely expressed, with higher expression in the liver, white adipocyte tissue (WAT) and pancreas (Fig. S6A and S6B).

Several studies have reported that starvation- or exercise-induced autophagy can lead to beneficial metabolic effects ^40, 53, 54^. After 36 hours of fasting, we found the phosphorylation of pS404-STK11IP level decreased (Fig. S6C), and STK11IP KO mice showed a robust increase of LC3-I to LC3-II conversion under these conditions (Fig. 4A and S6D). These results further indicate that STK11IP and its phosphorylation of pS404 could play a role in this process. This enhancement of autophagy significantly improved the fasting-induced fatty liver shown as the decrease of lipid droplet in the STK11IP knockout mice (Fig. 4B, 4C and 4D). There was also a concomitant decrease of TG and NEFA levels in serum and liver (Fig. 4E, 4F and 4G). STK11IP KO mice also exhibited relatively higher serum glucose levels compared to WT mice (Fig. 4H). These observations are consistent with the previous report that autophagy promotes the conversion of lipids to glucose during long-term fasting ^55^.

**Fig. 4.**
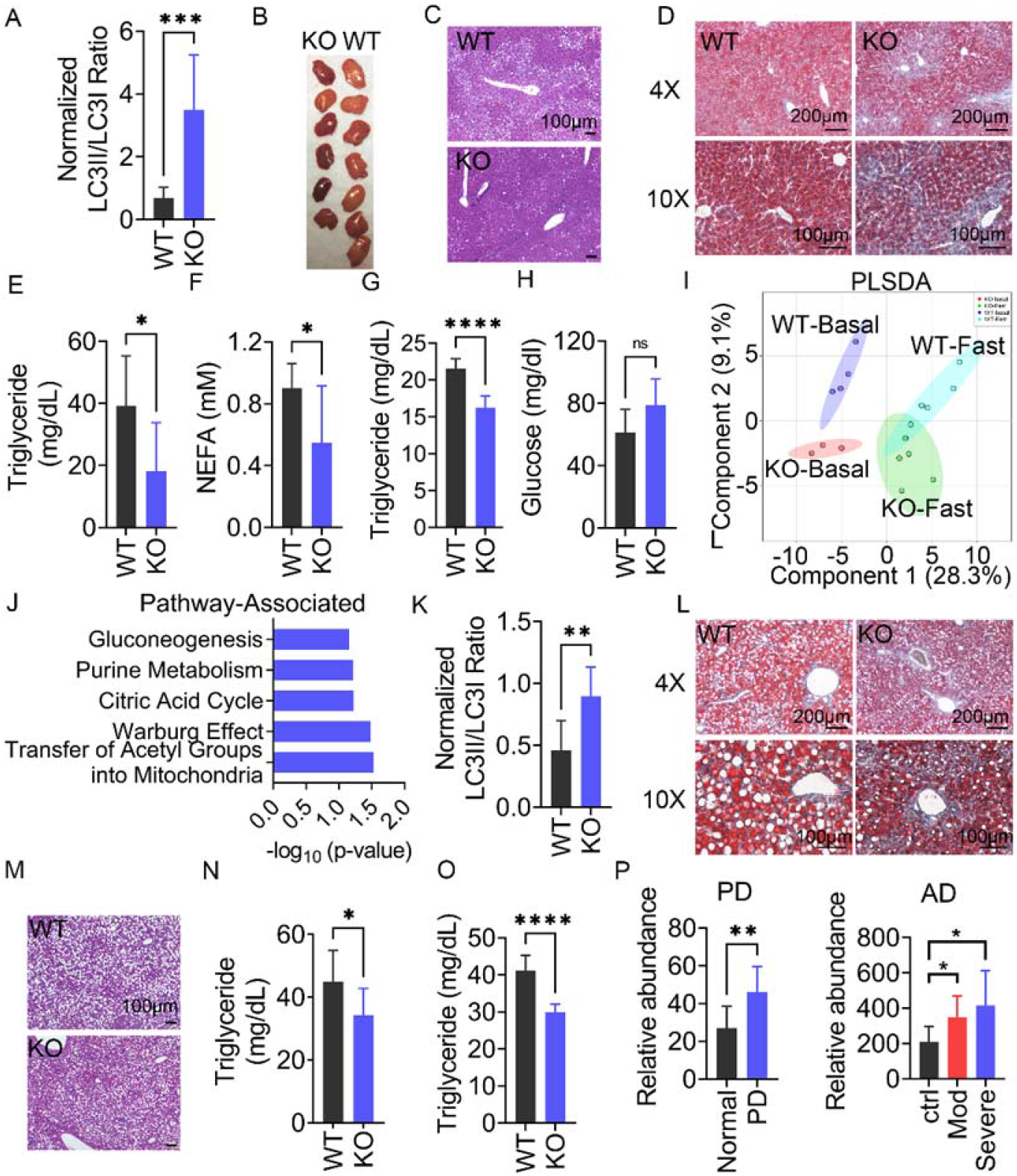
STK11IP Deficiency Protects Mice against Metabolic Disorder. (A) Statistics of the LC3II/LC3I ratio in WT and STK11IP knock out mice. The mice were subject to 36 hours of fasting. SE (short exposure); LE (long exposure); (WT, n=8; KO, n=6); t test. Results represent mean ± SEM. (B) Images of the liver from STK11IP WT (wild type) and KO (knock out) mice after 36 hours of fasting (WT, n=7; KO, n=6). (C-D) Histological sections (H&E stained) (C) and Oil-Red staining (D) of the liver from STK11IP WT (wild type) and KO (knock out) mice. The mice were subject to 36 hours of fasting after 2 months of HFD diet. (WT, n = 10; KO, n=9). Scales bars, 100 μm. (E-H) Blood triglycerides (E), NEFA (non-esterified fatty acids) (F), Liver triglycerides (G) and blood glucose (H) levels in wild type and STK11IP KO mice. Mice were subject to 36 hours of fasting (WT, n=8; KO, n=6); t test. Results represent mean ± SEM. (I) Partial least squares discriminant analysis (PLSDA) of the blood metabolites from WT (Wide-type) or KO (STK11IP-/-) mice (basal and 36 hours of fasting). (J) Pathway enrichment analyses of the serum metabolites that were significantly different between the WT and STK11IP KO mice after 36 hours fasting. The analysis was performed using the MetaboAnalyst 3.0 software. (K) Statistics of the LC3II/LC3I ratio in the liver from wild type and STK11IP knock out mice. The mice were subject to two weeks of MCD diet. SE (short exposure); LE (long exposure); (n = 6 per group); t test. Results represent mean ± SEM. (L-M) Histological sections (H&E stained) (M) and Oil-Red staining (L) of the liver from wild type and STK11IP KO mice. The mice were subject to 2 weeks of MCD diet. (WT, n = 8; KO, n=7). Scales bars, 100 μm. (N-O) Blood triglycerides (N) and Liver triglycerides (O) levels in STK11IP WT (wild type) or KO mice. The mice were subject to 2 weeks of MCD diet (WT, n = 8; KO, n=7); t test. Results represent mean ± SEM. (P) STK11IP Expression is Increased in Neurogenerative Diseases. STK11IP expression in control individuals, or patients with Parkinson’s disease (PD) or Alzheimer’s disease (AD). The data for the Parkinson’s disease patients and Alzheimer’s disease patients were extracted from the database with a reference number of GSM184355 and GSM697332, respectively; (normal: n=7, PD: n=17; n=7 for each group in AD), t test. Results represent mean ± SEM.

To gain further insights into the metabolic remodeling after STK11IP KO, we performed metabolomic profiling of the serum harvested from WT and STK11IP KO mice. Partial least squares discriminant analysis (PLSDA) showed distinct metabolic patterns for the WT and STK11IP KO groups under both basal and fasted conditions (Fig. 4I, S6E and table S6). Quantitative pathway enrichment analyses revealed a significant enrichment of glucogenesis, ketone body metabolism, beta-oxidation of fatty acids, and betaine or carnitine metabolism (Fig. 4J and S6F). These alterations may explain why the glucose level was higher and the TG level was lower in STK11IP KO mice after long-term starvation (Fig. 4E-4H). This is consistent with previous reports that upon starvation, fatty acids and amino acids generated by autophagy can be used to drive gluconeogenesis and ketogenesis. As starvation continues, degradation of adipose and muscle tissues plays an increasing role in supplying substrates to the liver, which exports glucose and ketone bodies to feed the other tissues ^55^.

Metabolic perturbations, including insulin resistance and altered lipid metabolism, have been hypothesized to contribute to the pathogenesis of NASH (nonalcoholic steatohepatitis), which is characterized by the imbalance in triglyceride (TG) production, uptake and removal in the liver. Mechanisms that promote fat clearance is expected to reduce NASH. We therefore utilized a NASH model that is induced by the Methionine-Choline-deficient (MCD) diet ^56–58^. After 2 weeks of the MCD diet, STK11IP KO mice showed a similar decrease of body weight, compared with WT mice (Fig. S6G). However, compared to control cells, the liver from STK11IP KO mice showed increased LC3II/LC3I ratios (Fig. 4K and S6H). Consistent with this, STK11IP KO mice showed reduced liver steatosis, as indicated by the reduced macro-vesicular lipid droplets (Fig. 4L and 4M) and the downregulation of TG levels in the serum and liver tissue (Fig. 4N and -4O).

## Discussion

mTORC1 functions at the convergence point of a vast signaling network that senses fluctuations in extracellular and intracellular nutrients to regulate a large variety of cellular anabolic processes ^59^. The activation of mTORC1 is mainly achieved by two conditions, the nutrient-dependent activation of the heterodimeric Rag-GTPases that recruit mTORC1 to the lysosomal surface, together with the growth factor-driven activation of the lysosome-bound Rheb-GTPase ^4^. Once mTORC1 is activated, whether mTORC1 phosphorylates local lysosomal proteins to mediate its downstream signaling output, however, is poorly understood. To identify the local lysosomal substrates of mTORC1, we combined and cross-referenced two datasets, i.e., a lysosomal proteomic dataset (Fig. 1A) and our previous hepatic mTORC1-regulated phosphoproteomic database ^16^. These integrative analyses led to identification of a poorly studied protein, STK11IP ^34^, as a potential lysosomal mTORC1 substrate (Fig. 1B and 1C). Further biochemical studies showed that: (1) STK11IP is localized on the lysosomal membrane, (2) the phosphorylation level of STK11IP at S404 correlates with the activity of mTORC1 in cells, (3) STK11IP interacts with Raptor, but not Rictor, and (4) STK11IP S404 is phosphorylated by mTOR *in vitro*. These results indicate that STK11IP is a novel, lysosome-localized substrate of mTORC1. In addition to STK11IP, our database also contains many lysosomal proteins, as well as other potential non-lysosomal mTORC1 substrates that will be useful for future research focused on lysosomal biology and mTORC1 signaling (Fig. 1B). Finally, the integration of organelle proteomic and global quantitative phosphoproteomic results offers a general strategy for the identification of the organelle-specific substrates for a kinase of interest.

Autophagy is a self-digestion process that is critically involved in the elimination of defective/accumulated proteins and damaged organelles, as well as removal of intracellular pathogens. The dysregulation of autophagy is implicated in the pathogenesis of a broad ranges of diseases, including aging, metabolic syndrome, neurodegenerative disease, cancer, and heart disease ^52^. Lysosomes, which are known as the cellular “incinerators”, not only provide the platform for mTORC1 activation, but receive signals from mTORC1 to regulate autophagy. Besides STK11IP, TFEB is the only known mTORC1 substrate that is also localized on the lysosome ^18, 19, 60^. Our data revealed a novel, lysosomal acidification-related mechanism by which mTORC1 regulates autophagy (Fig. 3O). We found that STK11IP phosphorylation could regulate V-ATPase activity through the interaction of STK11IP with ATP6V1A (mediated by the domain of 360-770 of STK11IP (Fig. S5E and S5F)). How STK11IP and V-ATPase interactions regulate the activity of V-ATPase, however, will be addressed by future structural studies of this protein complex. Finally, a recent study showed that the luminal pH values of individual lysosomes are unequal, and this heterogeneity could result from the position of each lysosome within the cell^61^. Our data thus raise the intriguing hypothesis whether the distribution of activated mTORC1 and hence pS404-STK11IP status could be a mechanism to explain the position-dependent regulation of lysosomal acidification.

We explored the *in vivo* implications of our findings using an STK11IP KO mouse model. During starvation, autophagy in the liver generates nucleosides, fatty acids and amino acids to provide the substrates for ketogenesis and glucogenesis. Inhibition of autophagy will result in more lipids storage in the liver ^55, 62^. In this study, we observed that STK11IP KO mice showed enhanced autophagy. Under starvation conditions, the serum metabolites that were differentially present between WT and STK11IP KO mice were mainly linked to ketone body metabolism and glucogenesis (Fig. 4J and S6F). As a result, STK11IP KO mice exhibited a lower serum TG and a higher blood glucose levels relative to their controls, which is consistent with enhanced autophagy in STK11IP KO mice (Fig. 4E-4H). In addition, metabolic perturbations, including insulin resistance and impaired lipid metabolism, have been hypothesized to contribute to the pathogenesis of NASH (nonalcoholic steatohepatitis). Autophagy is known to be deregulated during the pathogenesis and progression of NASH. Impaired autophagy prevents the clearance of excessive lipid droplets. Rapamycin treatment or FNDC5 KO results in enhanced autophagy, which can ameliorate NASH ^63, 64^. Indeed, we found that STK11IP KO can reduce MCD diet-induced NASH in mice (Fig. 4K-4O and S6H).

Because autophagy also plays a critical role in neurodegenerative diseases, we also searched public databases, and found that STK11IP expression is upregulated during the progression of neurodegenerative diseases, including the Parkinson’s disease and Alzheimer’s disease (Fig. 4P). Intriguingly, using a large-scale, multi-omics profiling approach, ATP6V1A (one of the STK11IP interacting proteins) has recently been identified as a key regulator of the pathogenesis of sporadic late-onset Alzheimer’s disease (LOAD) ^65^. All these results point to STK11IP as a critical player in the regulation of autophagy implicated in neurodegeneration. It is important to note that STK11IP depletion has no effect on the activity of mTORC1. Agents that pharmacologically target STK11IP therefore will offer a strategy to manipulate autophagy, avoiding the known side effects associated with mTORC1 inhibitors (e.g., rapamycin) ^66–70^. The full therapeutic potential of these agents in the treatment of the relevant diseases will be addressed in the future studies.

In summary, we used quantitative proteomics and identified STK11IP as a novel, lysosomal-specific substrate of mTORC1. STK11IP and its mTORC1-mediated phosphorylation regulates lysosome acidification and autolysosome maturation. This offers a novel, TFEB/ULK1-independent mechanism for mTORC1 to regulate autophagy. STK11IP ablation leads to enhanced autophagy in mice, which protects these animals against NASH. This study provides the foundation for targeting STK11IP for the treatment of diseases with aberrant autophagy signaling.

## Acknowledgements

We thank Dr. Joseph Albanesi for the helpful discussions. We thank Dr. Hamid Baniasadi for the help with metabolomic studies, Metabolomics Core Facility within the Biochemistry Department. We thank the UT Southwestern Mouse Metabolic Phenotyping Core and Carlos Castorena for the assistance with mice treadmill exercise studies. We also thank the University of Texas Southwestern Medical Center McDermott Center Next Generation Sequencing Core, Flow Cytometry Core of the Children’s Research Institute, Histopathology Core, Live Cell Imaging Core, and Animal Resource Center. This work was supported in part by grants from the Welch Foundation (I-1800 to Y.Y.), NIH (R35GM134883, R01GM122932 and R01GM114160 to Y.Y.), and CPRIT (RP160157 to Z.Z.).

## Materials and Methods

### Materials

All the antibodies, plasmid and reagents were listed in table S7.

### Cell lines

HEK293T (ATCC, Cat# CRL-3216) and primary MEF cells (isolated from wild type and STK11IP KO mice) were cultured in Dulbecco’s modified Eagle’s medium (DMEM, sigma, D5796) supplemented with 10% fetal bovine serum (FBS, sigma, 12306C) and antibiotics (Gibco, 15240112) at 37 □ in a 5% CO_2_ incubator.

HEK293T cells were transfected with the Cas9-sgRNA that targeted to STK11IP or GFP. Single cell clones were screened using flow cytometry, and were confirmed for the KO efficiency using immunoblotting experiments.

All the reconstituted cells (WT/S404A/S4-4D cells; GFP-LC3 cells, GFP-LC3-RFP cells and LAMP1-GFP-Flag cells) were screened single cells with the same expression level, measured using western blot.

HEK293TD cells were cultured to 50–70% confluency in 10 cm dishes, and were then transfected using Lipofectamin-2000 (Sigma). The Plenti or pLKO.1 plasmid, VSVG, and delta8.9 were co-transfected at 8 μg, 6 μg, and 4 μg, respectively. The medium was changed 6 hours after the transfection. Viruses were collected twice at 24 h and 48 h after the transfection. Subsequently, 1 mL of virus was added to each well of WT (wild type) and KO (STK11IP KO) HEK293T cells in 6-well plates with Polybrene at a final concentration of 8 μg ml^-1^. After splitting the cells once, cells were infected with virus again using the same procedure. The medium was replaced after 48 h with a fresh growth medium containing 2 μg ml^-1^ blasticidin or puromycin.

### LC3 turnover assay

WT and STK11IP KO HEK293T cells were cultured in regular DMEM with or without 2 hours of 20 μM chloroquine or 100 nM Bafilomycin A1 treatment. The cells were using the lysis buffer (1% SDS, 10 mM HEPES, pH 7.0, 2 mM MgCl_2_, 20 U ml^-1^ universal nuclease). Same amounts of proteins were subjected to immunoblotting analyses. Quantification of western blots was performed using Image J.

### Mice and treatment

Wild-type (005304), STK11IP KO mice (028999) and RFP-GFP-LC3 transgenic mice (027139) were purchased from the JAX lab.

### Genotyping

Mice were genotyped followed the protocol provided by the JAX lab. STK11IP KO mice were genotyped using primers (533: CAG TGT GCT ACA GCC AGA GAG; 532: GAG CTG GGG AGG AGG TAG AC; 534: AGG CCA TCT CTC TGT CCT CA). mRFP-GFP-LC3 mice were genotyped using primers (24935: CAT GGA CGA GCT GTA CAA GT; 24936: CAC CGT GAT CAG GTA CAA GGA; oIMR7338: CTA GGC CAC AGA ATT GAA AGA TCT; oIMR7339: GTA GGT GGA AAT TCT AGC ATC ATC C)

Mice were maintained on a 12-hour dark/12-hour light cycle and housed in groups of 3–5 with unlimited access to water and food. Normal chow diet was LabDiet (5058); high-fat diet (60% fat, D12451) and MCD diet (methionine and choline deficiency diet, A02082002BR) were purchased from Research Diet. The Institutional Animal Care and Use Committee of the University of Texas Southwestern Medical Center approved all animal experiments. All efforts were made to follow the Replacement, Refinement and Reduction guidelines. To minimize discomfort, mice were anesthetized with a ketamine/xylazine cocktail or 2%–3% isoflurane during surgery or before organ harvest. The STK11IP KO mice were backcrossed to 6^th^ generation with wild-type C57BL/6NJ mice.

### HFD or MCD diet feed

The littermate mice with similar body weight were feed with high fat diet (60% fat, D12451) from about 2-month-old, or MCD diet (methionine and choline deficiency diet, A02082002BR) from 2-month-old. Body weights were measured on a daily basis.

### TG, NEFA, Cholesterol measurement

Blood samples were collected from the tail vein. The same amount of liver tissue samples was collected and weight. The blood or liver triglycerides, blood cholesterol and NEFA levels were measured following the manufacture’s instruction.

### Isolation of the lysosomes

Confluent HEK293T cells that stably expressing LAMP1-GFP-FLAG^X3^ (LGF) or STK11IP-GFP-Flag^X3^ (SGF) were rinsed with cold PBS, scraped, spun down and resuspended in 750 μl of fractionation buffer: 140 mM KCl, 5 mM MgCl_2_, 50 mM Sucrose, 20 mM HEPES, pH 7.4, supplemented with protease inhibitors. The cells were mechanically broken with a pellet pestle and were spun down at 1000 g for 10 min to remove nucleus and cells debris. The samples were then spun down at 20,000 g for 25 min to yield the organelles pellets. The pellet was resuspended in fraction buffer and subjected to immunoprecipitation with 100 μl of a 50% slurry of anti-FLAG magnetic beads (M8823, Sigma). Four hours later, beads were washed 4 times using fractionation buffer, then eluted with 1% SDS lysis buffer (1% SDS, 10 mM HEPES, pH 7.0, 2 mM MgCl_2_, 20 U ml^-1^ universal nuclease), and the proteins were subjected to immunoblot or MS analysis.

For the isolation of lysosome using ultracentrifugation method was performed following the manufacturer’s instructions (Lysosome Isolation Kit, Sigma-Aldrich, LYSISO1-1KT).

### RNA extraction and qRT-PCR

Cells or tissues were lysed in the TRIzol reagent (Sigma, T9424), and mRNA was extracted according to the standard protocols. Reverse transcription was performed using the SuperScript™ III One-Step RT-PCR System (Invitrogen, 12574026) according to the manufacturer’s instructions. qRT–PCR was performed with a real-time system (SYBR green, Applied Biosystems, A25741). Data were normalized to the internal control and presented as relative expression levels.

### Co-Immunoprecipitation

HEK293T cells were rinsed once with ice-cold PBS 24-36 hours after the transfection and lysed in the ice-cold BLB lysis buffer (10 mM KPO_4_, pH 7.6, 6 mM EDTA, pH 8.0, 10 mM MgCl_2_, 0.5 % NP-40, 0.1 % BriJ-35, 0.1 % DOC, adjust pH to 7.4) or IP buffer (0.5% NP-40, 150 mM NaCl, 50 mM Tris-HCl, pH 7.5) added with EDTA-free protease inhibitors (Roche). The soluble fractions from cell lysates were isolated by centrifugation at 13,000 rpm for 10 minutes. For immunoprecipitations, 20 μl of a 50% slurry of anti-FLAG (A2220, Sigma) or anti-HA beads (A2095, Sigma) were added to each lysate and incubated with rotation for overnight at 4°C. Immunoprecipitants were washed three times with BLB lysis buffer. Immunoprecipitated proteins were denatured by the addition of 100 μl of sample buffer and boiling for 10 minutes and subjected to immunoblot analysis.

### Histology and Immune Staining

Tissues were fixed in 4% paraformaldehyde and were embedded in paraffin or frozen sections. Paraffin sections (5 μm) were deparaffinized and the frozen sections were washed with PBS to remove the OCT compound. The hematoxylin (Vector, H3401) and eosin Y (Thermo, 6766007) staining were performed according to standard protocols or the manufacturer’s instructions.

Cells were plated on micro cover glasses (Electron Microscopy Sciences 72226-01). After confluence, cells were washed with PBS and fixed with 4% paraformaldehyde (PFA) in PBS for 10 min. The cells were permeabilized (0.2% Triton X-100, 10 min), blocked in 3% BSA (Sigma, A9418), then incubated with primary antibodies overnight at 4°C (for example, the LC3, LAMP2 and Tom20 antibody). After the incubation with primary antibodies, slides were washed and incubated with Alexa Fluor 546 goat anti-rabbit or Alexa Fluor 488 goat anti-mouse secondary antibodies (Life Technologies, A11001 and A11010) at room temperature for 1 h. The slides were then washed and stained with DAPI (Sigma, D9542), and sealed with mounting medium (FluorSave reagent, Millipore, 345789). Photomicrographs were taken of representative 10×, 20× or 40× magnification fields using confocal microscope (Confocal Zeiss LSM880 Airyscan).

### Flow cytometry

Cells were washed, trypsinized, and centrifuged at 1,000 r.p.m. for 4 min. Cells were washed once with FACS buffer (DPBS with 2% FBS), then resuspended in FACS buffer for FACS analysis (LSRFortessa, BD) or sorting (FACS Aria II SORP, BD).

### Immunoblot analysis and Subcellular fraction

For immunoblot analysis, the cells were lysed using 1% SDS lysis buffer (1% SDS, 10 mM HEPES, pH 7.0, 2 mM MgCl_2_, 20 U ml^-1^ universal nuclease with added protease inhibitors and phosphatase inhibitors). Protein concentrations were measured using the BCA assay (23227, Thermo Fisher). The lysates were mixed with the 4X reducing buffer (60 mM Tris-HCl, pH 6.8, 25% glycerol, 2% SDS, 14.4 mM 2-mercaptoethanol, 0.1% bromophenol blue). Samples were boiled for 10 min and the same amounts of proteins were subjected to electrophoresis using the standard SDS-PAGE method.

Proteins were then transferred to a 0.22 μm nitrocellulose membrane (GE, 10600001), and were blocked with a TBST buffer (25 mM Tris-HCl, pH7.5, 150 mM NaCl, 0.05% Tween-20) containing 3% non-fat dried milk and incubated overnight with primary antibodies (1:1,000 dilution) at 4 °C and for 1h at room temperature with peroxidase-conjugated secondary antibodies (1:10000 dilution). Blots were developed using enhanced chemiluminescence, and were exposed on autoradiograph films and were developed using standard methods.

Segregation and enrichment of proteins from different cellular compartments were performed using a Thermo Scientific Subcellular Protein Fractionation Kit (78840, Thermo Fisher) followed the manufacturer’s instructions. The same amount of protein was subjected to immunoblot analysis. All the western blot images were quantified by using the software package ImageJ.

### Isolation of Primary MEF cells

Mouse embryonic fibroblasts (MEF) were isolated from wild-type and STK11IP KO mice or WT/STK11IP KO mice crossing with mRFP-GFP-LC3 mice following the previous protocol ^71^.

### Endurance exercise test

For the endurance exercise test, 4-month-old mice (wild type and STK11IP KO cell with or without mRFP-GFP-LC3) were acclimated to and trained on an exercise treadmill (Columbus Instruments) for 2 days. On day 1 mice ran for 5 min at 8 m min^-1^ and on day 2 mice ran for 5 min at 8 m min^-1^ followed by another 5 min at 10 m min^-1^. On day 3, mice were subjected to a single bout of running starting at the speed of 10 m min^-1^. Forty minutes later, the treadmill speed was increased at a rate of 1 m min^-1^ every 10 min for a total of 30 min, and then increased at the rate of 1 m min^-1^ every 5 min until mice were exhausted. Exhaustion was defined as the point at which mice spent more than 5 s on the electric shocker without attempting to resume running. Total running time was recorded, and total running distance was calculated for each mouse ^40, 72^

### Metabolomics

For metabolites extraction, ice-cold methanol was added to the blood samples to a final concentration at 80%. Metabolite profiling was performed with LC–MS/MS. The peak area for each detected metabolite was normalized to the total ion count. The preprocessed datasets were mean-centered, and unit-variance scaled. Principal component analysis and hierarchical clustering of metabolites in different samples were performed using MetaboAnalyst 3.0. ^73^.

### Biotin labeling with TurboID in HEK293T cells

The TurboID-STK11IP stably expressing HEK293T cells were cultured in no-Biotin DMEM medium for five generations. Biotin (50 μM) labeling was then initiated with complete or amino acid depleted medium for 2 hours. The labeling was stopped by washing the cells with ice-cold PBS and the cells were lysed with the 1% SDS lysis buffer. The lysates were diluted to 0.1% SDS with PBS, and the proteins were immunoprecipitated using streptavidin beads. After the pull down, beads were washed for three times, and were boiled for 30 minutes. The eluate was subjected to immunoblot. For the MS sample, which was prepared using on-beads digestion following the previous report ^51^.

### Mass spectrometry experiment

Protein lysates were reduced by 2 mM dithiothreitol, and alkylated by adding iodoacetamide to a final concentration of 50 mM, followed by incubation in the dark for 20 min. The lysates were extracted by methanol-chloroform-precipitation and the proteins were solubilized in 8 M urea, then digested with Lys-C. The samples were then diluted to a final concentration of 2 M urea by the addition of 100 mM ammonium bicarbonate (pH 7.8). Proteins were digested overnight with sequencing-grade trypsin at a 1:100 (enzyme/substrate) ratio. Digestion was quenched by the addition of trifluoroacetic acid to a final concentration of 0.1%. For TMT MS samples, peptides were desalted using SepPak C18 columns (Waters) according to the manufacturer’s instructions. Peptides were fractionated by using an off-line two-dimensional SCX-RP-HPLC (strong-cation-exchange reversed-phase HPLC) protocol.

The MS samples were analyzed by LC-MS/MS experiments on an LTQ Velos Pro Orbitrap mass spectrometer (Thermo) using a top-20 CID (collision-induced dissociation) method. MS/MS spectra were searched against a composite database of the human protein database and its reversed complement using the Sequest algorithm. Search parameters allowed for a static modification of 57.02146 Da for cysteine and a dynamic modification of oxidation (15.99491 Da) on methionine, respectively. Search results were filtered to include <1% matches to the reverse database by the linear discriminator function using parameters including Xcorr, dCN, missed cleavage, charge state (exclude 1C peptides), mass accuracy, peptide length and fraction of ions matched to MS/MS spectra. Peptide quantification was performed by using the Core Quant algorithm ^74^.

For the data process of lysosome proteomics, we used the lysosome immunoprecipitation-proteomics experiments to define the lysosomal proteome. For these analyses, we performed two biological replicate experiments. After the MS experiments, we performed cross-reference analyses and excluded the proteins that were commonly identified in both the control (GFP-Flag) and the LGF (LAMP1-GFP-Flag) sample. These excluded proteins were classified as non-specific proteins. After filtering out these non-specifically bound proteins, we were able to derive a list of proteins that were specifically identified in the LGF sample. To increase the confidence in the protein identification, we also only selected the proteins that were identified with at least two unique peptides. To ensure that we identify the potential lysosomal proteins in an unbiased manner.

### LysoSensor/LysoTracker staining and pH measurement

Wild type/STK11IP KO cells or WT/S404A/S404D reconstituted cells were plated in 8 well charmers containing 200 μL complete DMEM medium. In a typical procedure, the cells were cultured in complete medium, then 25 μg/mL of TMR-conjugated hybrid nanoprobes were added and kept for 15 min at 37 °C. The medium was removed and washed. Thereafter, cells were incubated with complete medium for 1 hour at 37 °C. The TRITC (590 ± 25 nm) filters were used for TMR imaging.

For LysoTracker or LysoSensor imaging, the cells were incubated in complete DMEM medium, then LysoTracker^®^ Red DND-99 (100 nM) and LysoSensor (2μM) was added for one hour, then the fluorescence was monitored using confocal.

For pH value measurement, quantification of lysosomal pH was performed using a ratio-metric lysosomal pH dye LysoSensor Yellow/Blue DND-160. The pH calibration curve was generated as described previously^75^. Briefly, cells were labeled with 2 μM Lyso-Sensor Yellow/Blue DND-160 for 30 min at 37 °C in regular medium, and excessive dye was washed out using PBS. The labeled cells were treated for 10 min with 10 μM monensin and 10 μM nigericin in 25 m M MES calibration buffer, pH 3.5, 4.0, 4.5 5.0, 6.0 and 7.0, containing 5 m M NaCl, 115 m M KCl, and 1.2 m M MgSO_4_. Quantitative comparisons were performed in a 96-well plate, and the fluorescence was measured with a microplate reader at 37 °C. Light emitted at 440 and 535 nm in response to excitation at 340 and 380 nm was measured, respectively. The ratio of light emitted at 440 and 535 was plotted against the pH values in MES buffer, and the pH calibration curve for the fluorescence probe was generated from the plot and the result was analyzed using GraphPad Prism 9.

### V-ATPase activity assay

The lysosomes of wild type (WT)/STK11IP KO (KO) or WT/S404A/ S404D reconstituted HEK293T cells were loaded with LysoSensor™ Yellow/Blue DND-160, and performed following published procedures ^50^. The cells were washed with a buffer (90 mM K-gluconate, 50 mM KCl, 1mM EGTA, 20 mM Hepes, 50 mM sucrose, and protease inhibitor cock-tail, pH 7.3) before the treatment with 1 μM FCCP for 15 min in order to completely dissipate the lysosomal pH gradient. The FCCP-treated cells were harvested and homogenized using a Dounce homogenizer. To prevent lysosomal re-acidification during the course of fractionation, the cells were maintained in the continued presence of FCCP at 4°C, until the final wash of the isolated organelles. The organelles were resuspended in 1 ml buffer without FCCP. The fluorescence was monitored using a plate reader (emission at 440 and 535 nm in response to excitation at 340 and 380 nm, respectively). After the baseline determination (4 min), 5 mM MgCl2 and 5 mM ATP were added to the suspension, and fluorescence was monitored for 20 min. The ratio of light emitted at 440/535 were calculated and result was analyzed using GraphPad Prism 9.

### In vitro kinase assay

STK11IP was phosphorylated by mTOR in vitro. HEK293T cells that stably express Flag-STK11IP were treated with Torin1 (2 μM) for 6 hrs, and Flag-Tagged STK11IP was pulled down from these cells. Recombinant mTOR was purchased from Millipore (14-770) and recombinant 4EBP1 was purchased from Millipore (516678). Flag-STK11IP or 4EBP1 was incubated with or without mTOR in 50 μl reaction mixtures at 37°C for 60 mins. The reaction mixture contained 1X mTOR kinase buffer (Invitrogen), 1X protease inhibitor cocktail (Roche), 2 mM DTT, 10 μM ATP, 1 μg Flag-STK11IP, and 250 ng mTOR. Reaction was stopped by adding 16 μl of 4X SDS sample buffer. The samples were boiled at 95°C for 5 mins, and were then subjected to immunoblot analysis ^14^.

---

**Fig. S1.**
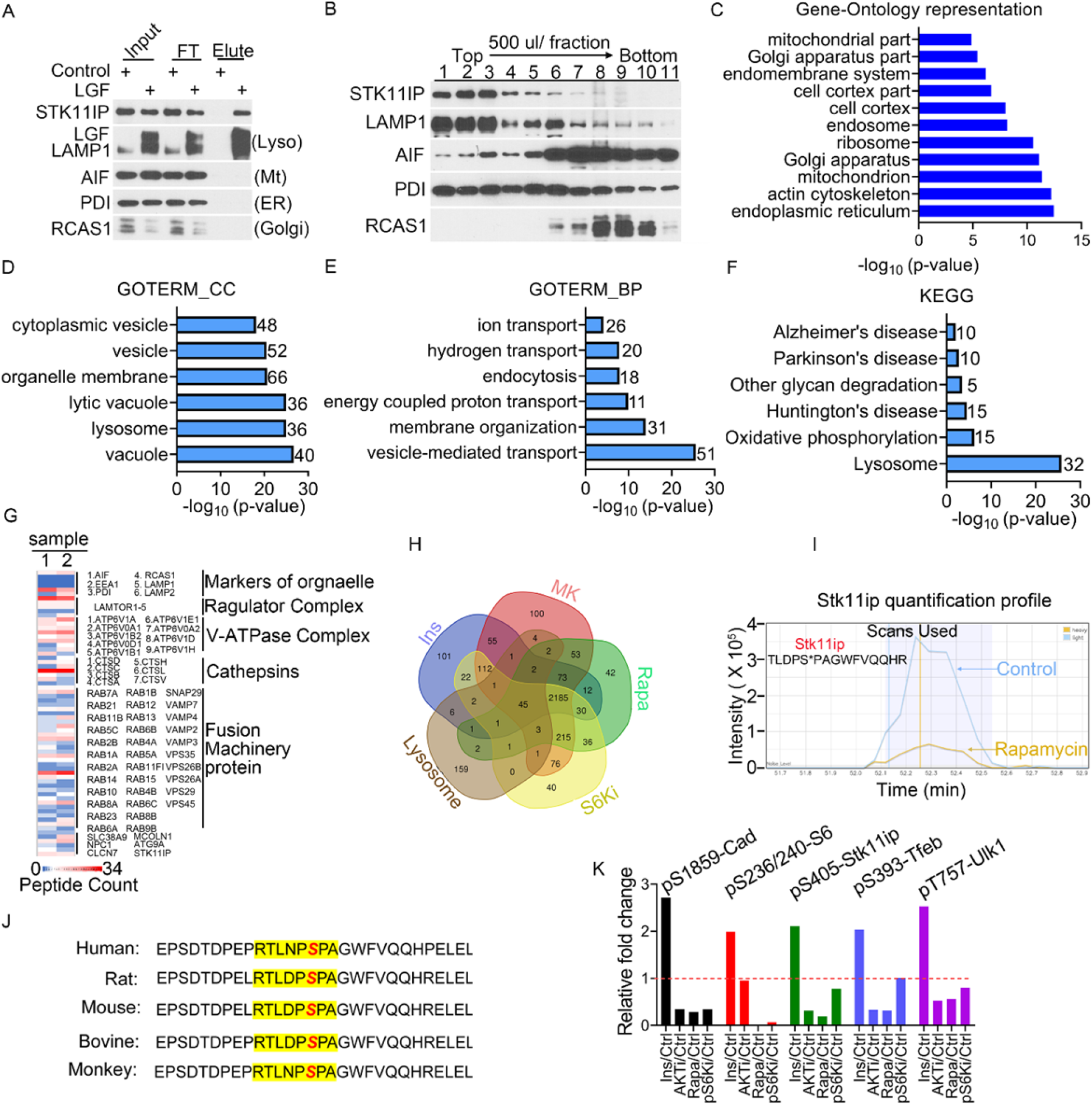
Integrative analyses of the mTORC1-regulated lysosomal phosphoproteome. (A) Immunoblot analyses of the immunopurified lysosomes. HEK293T cells stably expressing LAMP1-GFP-FLAG^X3^ (LGF) were extracted the organelles and subjected to immunoprecipitation against FLAG. The endogenous markers for several organelles were used, including LAMP1 (a lysosome marker, Lyso), AIF (a mitochondria marker, Mt), PDI (a marker for the endoplasmic reticulum, ER) and RCAS1 (a Golgi Complex marker, Golgi). (B) Immunoblot analysis of ultracentrifugation-purified lysosomes. The markers for the various organelles were the same as (A). (C) GO analyses of the proteins that are present in the ultracentrifugation-derived lysosomes, but not immunoprecipitation-derived lysosomes. (D-F) Gene Ontology (GO) analyses of the proteins identified in lysosome proteomic experiment. The following enrichment studies were performed: biological processes (BP) (D), cell component (CC) (E) and the KEGG pathway (F). (G) The relative abundances of the known lysosome proteins (data from two biological replicate experiments). (H) A Venn diagram demonstrating the proteins that were commonly identified between the hepatic phosphoproteomic experiments and the lysosomal proteomic experiments. Ins (Insulin); MK (MK2206, Akt inhibitor); Rapa (rapamycin, mTORC1 inhibitor); S6Ki (S6 kinase inhibitor). (I) The extracted ion chromatogram of the light (control, blue) and heavy (rapamycin-treated, yellow) ions of a Stk11ip peptide in the rat protein (TLDPS*PAGWFVQQHR, * indicates the phosphorylation site). (J) pS404-STK11IP is evolutionarily conserved among the various species. The S404 (in human) site is highlighted in red. (K) The level of changes of several representative phosphopeptides upon indicated condition, including pS1859-Cad, pS236/240-Rps6, pS405-Stk11ip (pS404 in the human proteins), pS393-Tfeb and pT757-Ulk1. Y=1 was labeled as red line. The data were extracted from the hepatic phosphoproteomic.

**Fig. S2.**
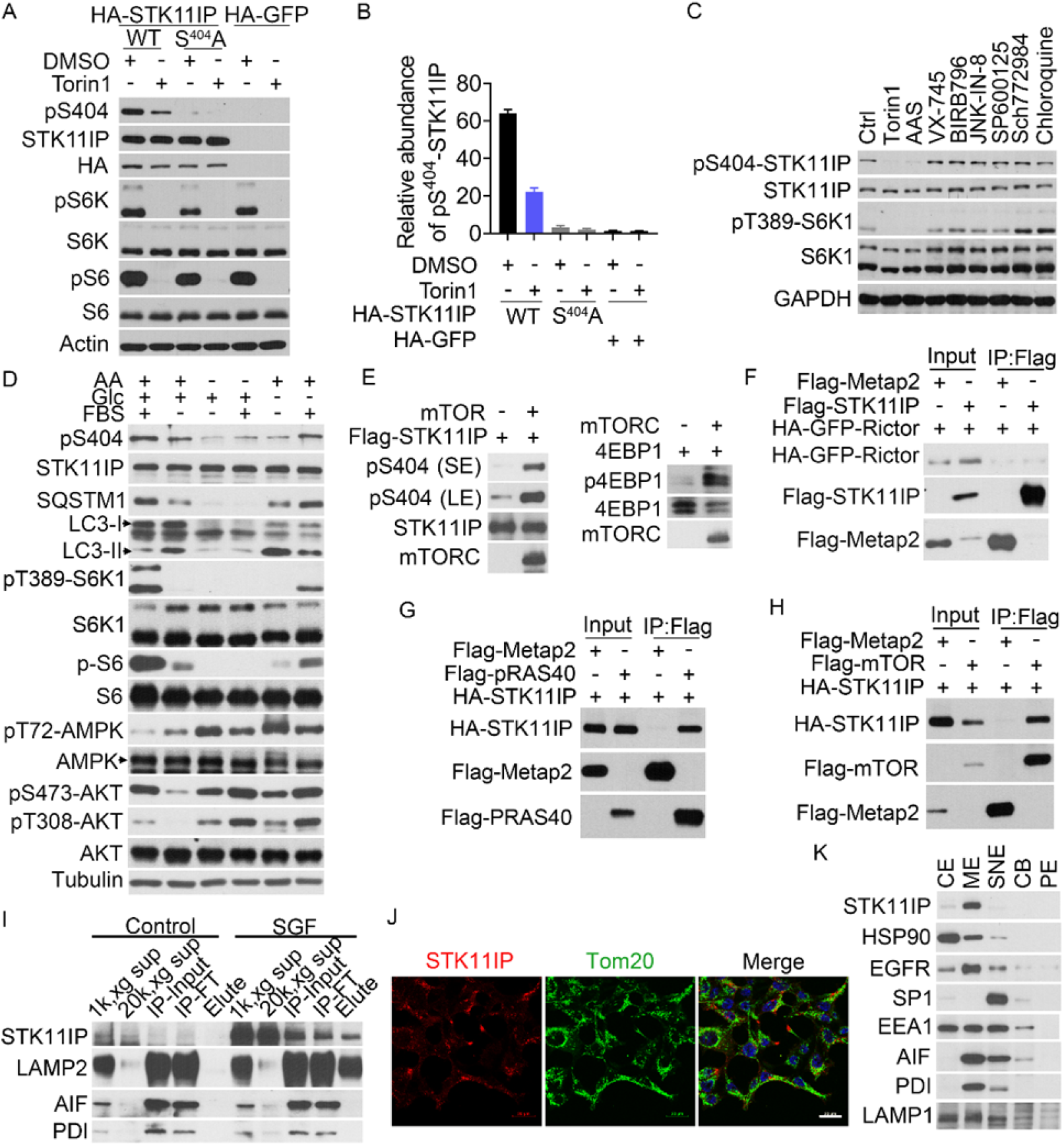
STK11IP is a lysosome-specific mTORC1 substrate. (A-B) The antibody against to pS404-STK11IP was specific to recognize pS404-STK11IP. Immunoblot analysis (A) of pS404, pS6K and pS6 in the HA-GFP, HA-WT-STK11IP and HA-S404A-STK11IP reconstituted cells upon DMSO (Ctrl) and Torin1 (2 μM for 4 hours) treatment. Quantification of pS404 level (B) for (A) was performed using Image J. (C) Immunoblot analysis of pS404, mTORC1 signaling in HEK293T cells upon indicated treatment for 4 hours. Ctrl (control, treat with DMSO), AAS (amino acid starvation), Torin1, Rapa (rapamycin); S6Ki (S6 kinase inhibitor); VX-745 (p38MAPK inhibitor), BIRB796 (p38/MAPK inhibitor), JNK-IN-8 (JNK inhibitor), SP600125 (JNK inhibitor) and Sch772984 (ERK1/2 inhibitor), CQ (chloroquine). (D) Phosphorylation of STK11IP at S404 is blocked as a result of nutrient deprivation. HEK293T cells were starved for the indicated nutrients (overnight by the indicated conditions). (E) STK11IP is phosphorylated by mTOR in vitro. Flag-STK11IP was prepared from cells (upon 6 hours Torin1 treatment) and incubated with recombinant mTOR in vitro. Phosphorylation of STK11IP at S404 was detected by using the phospho-specific antibody to this site, 4EBP1 as the positive control. SE (short exposer); LE (long exposure). (F-H) STK11IP does not interact with Rictor (F), but interacts with PRAS40 (G) and mTOR (H). Flag-tagged STK11IP or Metap2 (as the control) were co-transfected with HA-GFP-Rictor; or Flag-tagged PRAS40, mTOR and Metap2 (as the control) were co-transfected with HA-STK11IP into HEK293T cells. The lysates were subject to immunoprecipitation using anti-Flag antibody conjugated beads. (I) Immunoblot analysis of the various organelle markers from STK11IP-GFP-Flag (SGF)-immunopurified lysosomes. HEK293T cells stably expressing SGF were harvested, homogenized and subjected to immunoprecipitation against Flag. (J) Co-immunofluorescence analyses of STK11IP and Tom20 (a mitochondrial marker) in HEK293T cells. Scales bars, 20 μm. (K) STK11IP expression in different subcellular fractions. HEK293T cells were subject to subcellular fractionation, and each fraction was analyzed by immunoblot assays. HSP90, a cytoplasmic marker (CE, cytosol fraction); EGFR, a marker of the membrane (ME, membrane fraction); SP1, a marker for the soluble nuclear fraction (SNE); EEA1, a maker for the early endosome, CB (chromatin binding fraction), PE (pellet fraction).

**Fig. S3.**
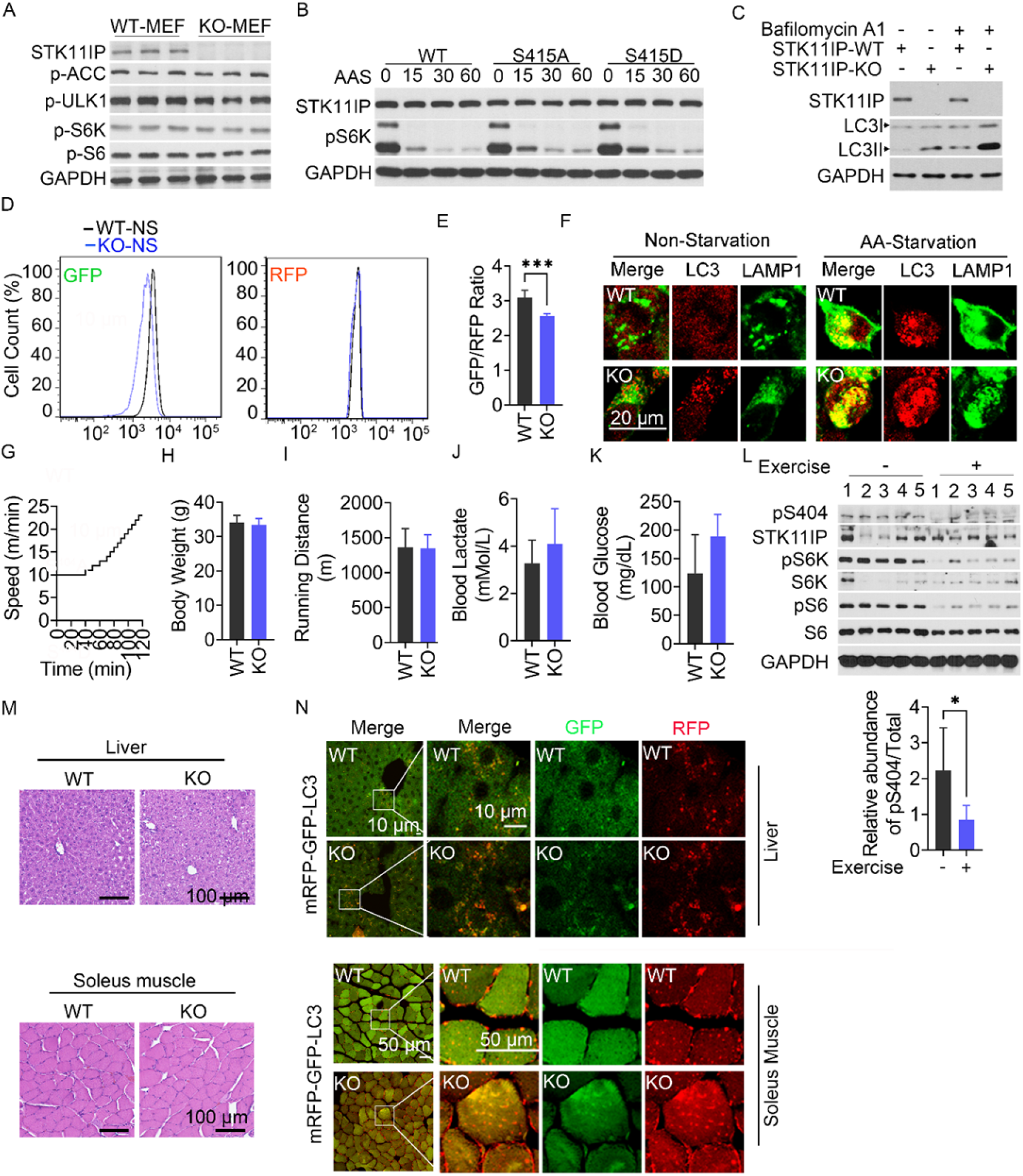
STK11IP Deficiency Promotes Autophagy. (A) STK11IP KO does not affect mTORC1 signaling. STK11IP WT (wild type) and KO primary MEF cells were lysed, and pS79-ACC, pS757-ULK1, pT389-S6K and pS240/244-S6 levels were probed using the indicated antibodies. (B) WT, S404A and S404D reconstituted HEK293T cells were amino acid starved (AAS) for 15, 30, 60 mins and subjected to immunoblot analyses using the indicated antibodies. (C) WT and STK11IP KO cells were cultured in complete media. When indicated, Bafilomycin A1 treatment was performed 100 nM for 2 hours. (D-E) GFP-LC3-RFP stably expressing WT and STK11IP KO cells were analyzed by flow cytometry (D) or plate reader (E) under normal medium; t test. Results represent mean ± SEM. (F) STK11IP knock out leads to more co-localization of LC3 and LAMP1, which indicate enhanced formation of autolysosome in STK11IP KO cells under non-starvation (complete media) and AA-Starvation (all amino acid deprivation) for 2 hours. Co-immunostaining of LAMP1 (Green signal) and LC3 (Red signal) was performed using STK11IP wild-type and knock-out cells. The cells were cultured in complete media. Scales bars, 20 μm. (G-K) Schematic showing the exercise regimen used in the endurance exercise test (G). WT and STK11IP KO mice were subject to this test. Afterwards, the body weight (H), treadmill running distance (I), blood lactate (J), glucose (K) of the WT (wild type) and STK11IP KO (KO) mice were measured (n=6 per group). Results represent mean ± SEM. (L) The pS404-STK11IP level decreased upon endurance treadmill exercise test. Biochemical analysis of pS404-STK11IP in Soleus Muscle from WT mice at rest (-) or after endurance exercise test (+) by Western blot detection (upper). GAPDH is shown as a loading control. Graphs showing quantification of relative ratio pS404/Total-STK11IP at rest and after exercise for 5 mice per group (bottom). Results represent mean ± SEM. (M-N) Histological sections (H&E stained) (M, Upper: liver; bottom: soleus muscle) and representative images of the mRFP-GFP-LC3 puncta (N, Upper: liver; bottom: soleus muscle) from the WT and STK11IP KO mice after the endurance treadmill exercise. (n = 6 per group). Scales bars, 100 μm (M); Scales bars, 10 μm (N, Upper: liver), 50 μm (N, Bottom: Soleus Muscle).

**Fig. S4.**
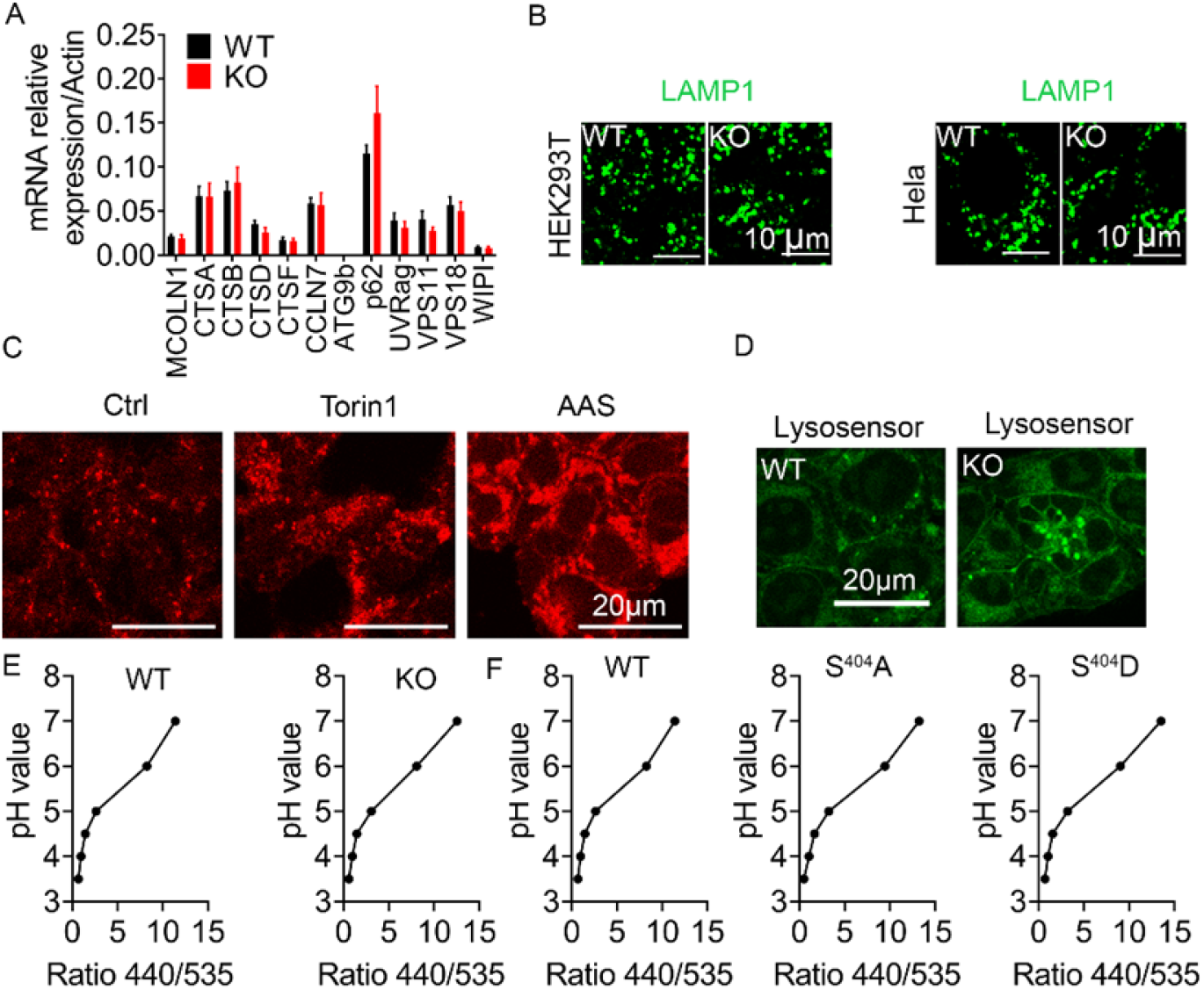
STK11IP Regulate Lysosomal Acidification. (A) STK11IP KO does not affect the expression of lysosomal genes. RT-qPCR analyses of representative autophagic and TFEB-target genes in WT and STK11IP KO primary MEFs. (n=3, per group); t test. Results represent mean ± SEM. (B) The lysosome morphology showed no difference between wild type and STK11IP knock out HEK293T or Hela cells. Immunostaining of LAMP1 was performed in the WT/KO-HEK293T and Hela cells. Scales bar, 10 μm. (C). Inactivation of mTORC1 by Torin1 (2 μM for 2 hours) or amino acid starvation (2 hours) lead to increased lysosomal acidity. Wild type HEK293T cells were stained with the Lysotracker dye upon indicated condition and subject to immunofluorescence analyses. Scales bar, 20 μm. (D) STK11IP knock out leads to increased lysosomal acidity. WT or STK11IP KO cells were stained with the Lysosensor dye and subject to immunofluorescence analyses. Scales bar, 20 μm. (E-F) The standard curve for pH measurement using the LysoSensor Yellow/Blue DND-160 dye In the Wild type and STK11IP KO cells (E) and WT, S404A or S404D reconstituted cells (F). These cells were subject to pH measurement using the LysoSensor Yellow/Blue DND-160 dye. (n=4 per point), Results represent mean ± SEM.

**Fig. S5.**
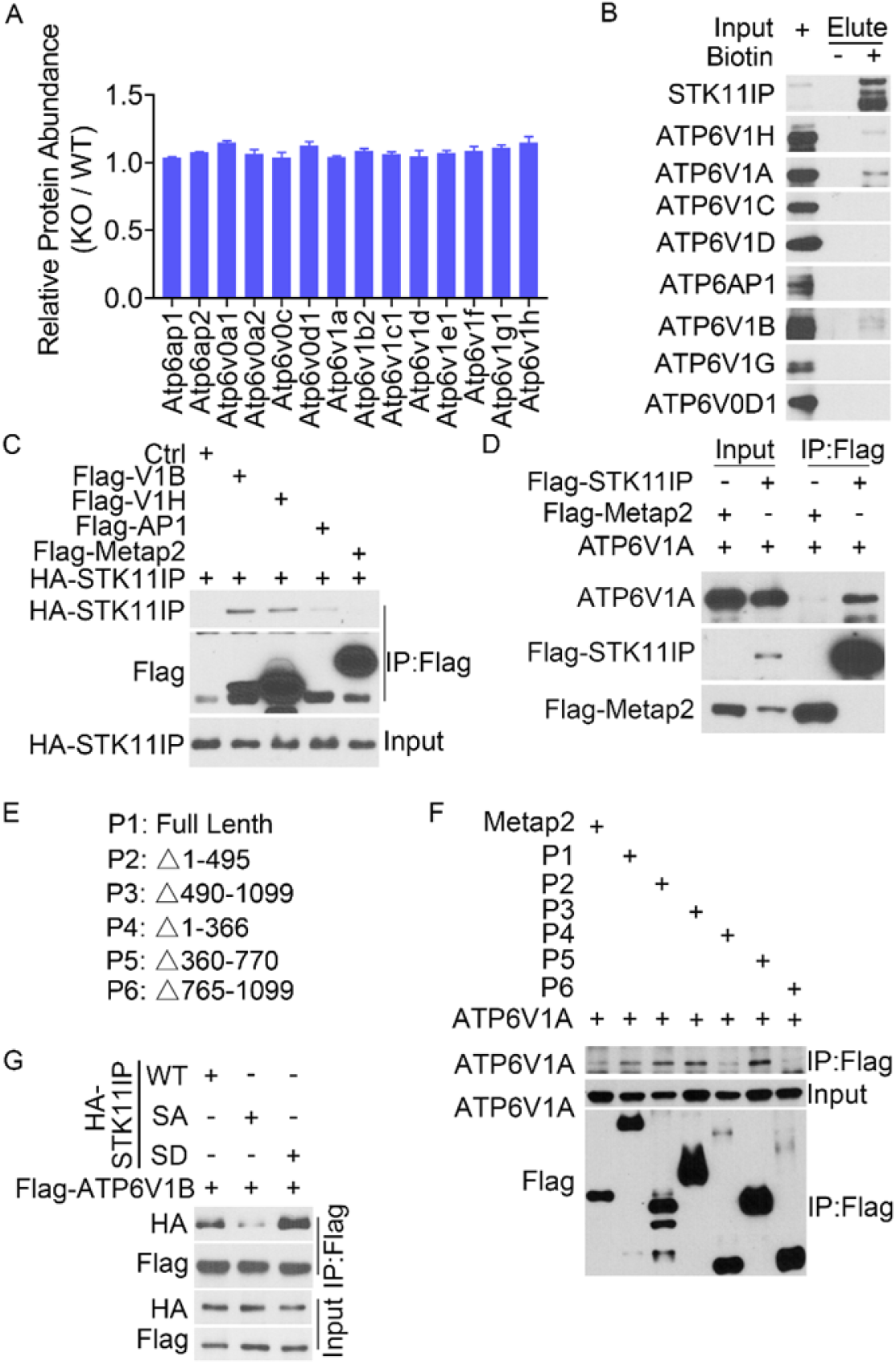
STK11IP Binds to V-ATPase complex. (A) Expression of the various lysosomal proteins as measured by TMT (tandem mass tag)-based quantitative proteomic experiments. WT and STK11IP KO primary MEF cells were subject to quantitative protein expression profiling experiments. The fold change (KO/WT) of protein abundances of the indicated proteins were extracted and presented. (n=3, for WT and KO cells), Results represent mean ± SEM. (B) Validation of the interaction between STK11IP and V-ATPase. TurboID-STK11IP stable cells were incubated with or without Biotin for 2 hrs. Cell lysates were subject to Streptavidin-based affinity purification. (C-D) STK11IP interacts with ATP6V1A, ATP6V1B and ATP6V1H. Flag or HA-tagged STK11IP was co-transfected with ATP6V1A, ATP6V1B-Flag or ATP6V1H-Flag into HEK293T cells. The lysates were subjected to immunoprecipitation. (E-F) Mapping the interaction domain of STK11IP with ATP6V1A. Flag tagged STK11IP truncation mutants (E) were co-transfected with ATP6V1A. The lysates were subjected to immunoprecipitation. (G) The STK11IP S404A (SA) mutant interacts more weakly with ATP6V1B, compared to STK11IP WT or the S404D (SD) mutant. HA-tagged STK11IP was co-transfected with ATP6V1B-Flag into HEK293T cells. The lysates were subjected to immunoprecipitation.

**Fig. S6.**
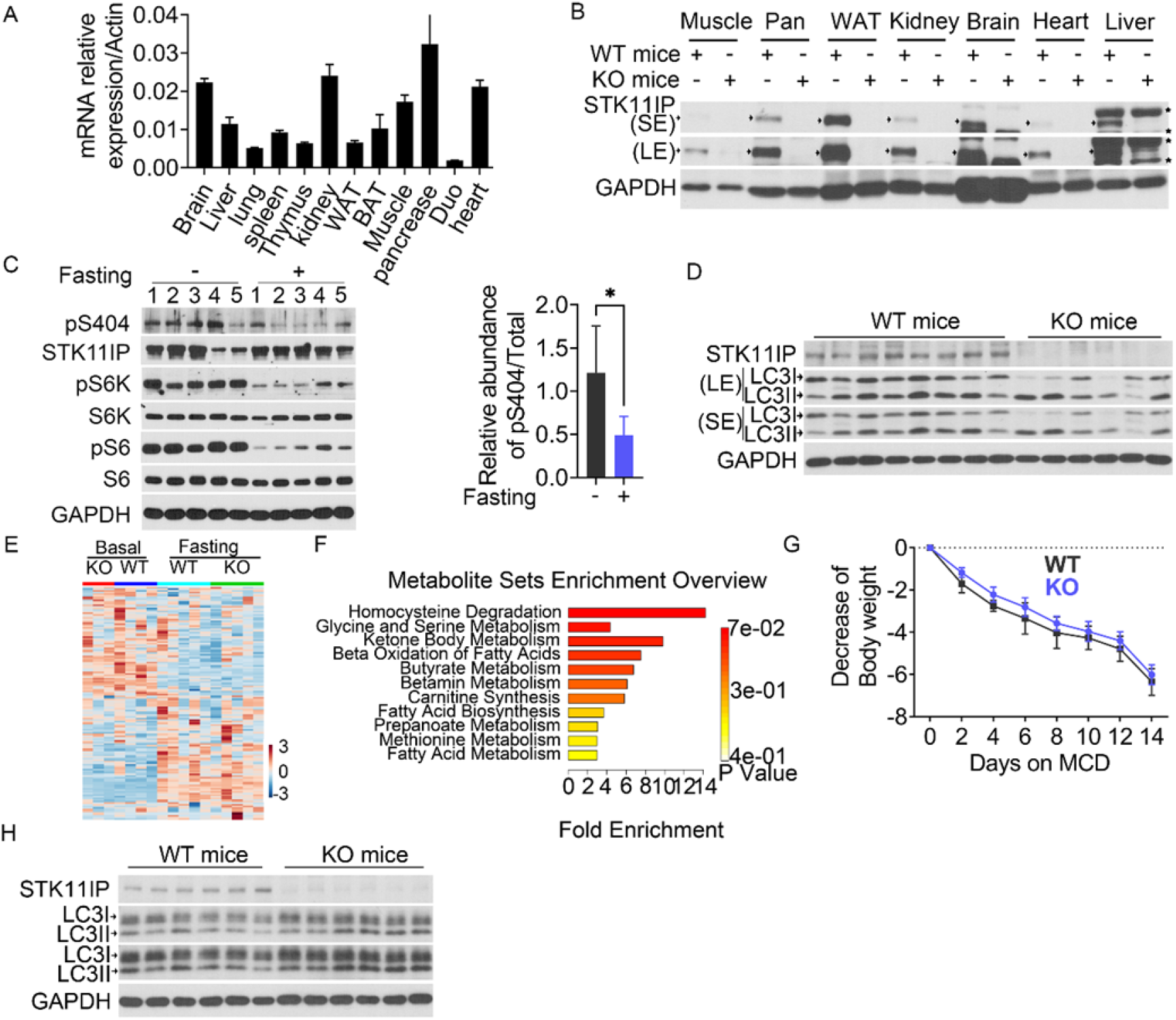
STK11IP Deficiency Protects Mice against Metabolic Disorder. (A-B) Expression levels of STK11IP in various mouse tissues, as measured by RT-qPCR analyses (A) and immunoblot analyses (B). The following tissues were harvested from 2-month-old mice, including the brain, liver, lung, spleen, thymus, kidney, white adipocyte tissue (WAT), brown adipocyte tissue (BAT), muscle, pancreas (Pan), duodenum (Duo) and heart. SE (short exposure); LE (long exposure). (n=3); t test. Results represent mean ± SEM. Arrow indicates STK11IP band, and star indicated non-specific band. (C) The pS404-STK11IP level decreased upon 36 hours of fasting. Biochemical analysis of pS404-STK11IP in liver from WT mice at normal (-) or after 36 hours fasting (+) by Western blot detection (Right). GAPDH is shown as a loading control. Graphs showing quantification of relative ratio pS404/Total-STK11IP at rest and after exercise for 5 mice per group (Left). Results represent mean ± SEM. (D) Immunoblot analysis the LC3II/LC3I ratio in WT and STK11IP KO mice. The mice were subject to 36 hours of fasting. SE (short exposure); LE (long exposure); (WT, n=8; KO, n=6). (E) The differentially expressed metabolites were subject to metabolic enrichment analyses using the MetaboAnalyst 3.0 software (WT-basal, n=4; WT-Fasting, n=5; KO-basal, n=3; KO-fasting, n=5). (F) Pathway enrichment analyses of the serum metabolites that were significantly different between the WT and STK11IP KO mice after 36 hours fasting (WT-basal, n=4; WT-Fasting, n=5; KO-basal, n=3; KO-fasting, n=5). (G) Body weight (in grams) of the WT and STK11IP KO mice (8 weeks old) that were fed with the MCD diet for 2 weeks (n=8 per group). Results represent mean ± SEM. (H) Immunoblot analysis LC3II/LC3I ratio in the liver from wild type and STK11IP-/- mice. The mice were subject to two weeks of MCD diet. SE (short exposure); LE (long exposure); (n = 6 per group).

